# An epitranscriptomic switch at the 5′-UTR controls genome selection during HIV-1 genomic RNA packaging

**DOI:** 10.1101/676031

**Authors:** Camila Pereira-Montecinos, Daniela Toro-Ascuy, Cecilia Rojas-Fuentes, Sebastián Riquelme-Barrios, Bárbara Rojas-Araya, Francisco García-de-Gracia, Paulina Aguilera-Cortés, Catarina Ananías-Sáez, Grégoire de Bisschop, Jonás Chaniderman, Mónica L. Acevedo, Bruno Sargueil, Fernando Valiente-Echeverría, Ricardo Soto-Rifo

## Abstract

During retroviral replication, the full-length RNA serves both as mRNA and genomic RNA (gRNA). While the simple retrovirus MLV segregates its full-length RNA into two functional populations, the HIV-1 full-length RNA was proposed to exist as a single population used indistinctly for protein synthesis or packaging. However, the mechanisms by which the HIV-1 Gag protein selects the two RNA molecules that will be packaged into nascent virions remain poorly understood. Here, we demonstrate that HIV-1 full-length RNA packaging is regulated through an epitranscriptomic switch requiring demethylation of two conserved adenosine residues present within the 5′-UTR. As such, while m^6^A deposition by METTL3/METTL14 onto the full-length RNA was associated with increased Gag synthesis and reduced packaging, FTO-mediated demethylation was required for the incorporation of the full-length RNA into viral particles. Interestingly, HIV-1 Gag associates with the RNA demethylase FTO in the nucleus and drives full-length RNA demethylation. Finally, the specific inhibition of the FTO RNA demethylase activity suppressed HIV-1 full-length RNA packaging. Together, our data propose a novel epitranscriptomic mechanism allowing the selection of the full-length RNA molecules that will be used as viral genomes.

## INTRODUCTION

Retroviral full-length RNA plays two key functions in the cytoplasm of infected cells. First, it is used as the mRNA template for the synthesis of Gag and Gag-Pol precursors and, second, it serves as the genomic RNA (gRNA) packaged into newly produced viral particles^1–3^. In contrast to the simple retrovirus murine leukemia virus (MLV), which was shown to segregate its full-length RNA into two functionally different populations serving as template for translation (mRNA) or packaging (gRNA), the HIV-1 and HIV-2 full-length RNA were proposed to exist as a single population acting indistinctly as mRNA and gRNA^4–6^. However and despite several years of efforts, there is still an important gap in our knowledge regarding the molecular mechanisms behind the selection of the full-length RNA molecules that will be incorporated into assembling viral particles.

The 5′-untranslated region (5′-UTR) present within the HIV-1 full-length RNA is the most conserved region of the viral genome and contains several high order structural motifs involved in different steps of the viral replication cycle from transcription, reverse transcription, splicing, translation to dimerization and packaging^3, 7, 8^. Since the full-length RNA serves both as mRNA and gRNA, translation and packaging are expected to be mutually exclusive events^2^. The Gag protein recognizes *cis*-acting RNA elements present at the 5′-UTR and the beginning of the Gag coding region and drives the selective incorporation of two copies of the gRNA into assembling viral particles. Indeed, there is accumulating evidence showing that dimerization and packaging of the HIV-1 full-length RNA are two tightly interconnected processes dependent on the Gag protein^9–11^. Structural and mutational analyses proposed that a conformational switch within the 5′-UTR regulates the transition from translation to dimerization and packaging *in vitro*^12–15^. In such models, the 5′-UTR alternates in conformations that occlude the dimerization initiation signal (DIS) or the Gag start codon thus, favoring translation or dimerization and packaging, respectively^3, 8^. However, chemical probing performed in cells and purified viral particles showed that a single structure, in which DIS is accessible for dimerization and packaging, predominates in these biological states^16, 17^ Moreover, the packaging prone structure does not interfere with full-length RNA translation suggesting that other factors rather than structural rearrangements are involved in the regulation of the cytoplasmic sorting of the HIV-1 full-length RNA^18, 19^.

It was recently reported that the HIV-1 full-length RNA contains N^6^-methyladenosine (m^6^A) residues located at the 5′- and the 3′-UTR as well as at internal positions such as the Rev response element (RRE)^20–22^. Methylation of adenosines at the RRE and the 3′-UTR was shown to promote Gag synthesis by favoring nuclear export and/or the intracellular accumulation of viral transcripts at late stages of the replication cycle^20–23^. However, it was also reported that the presence of m^6^A could also induce the degradation of the incoming gRNA early during infection^22, 23^. These controversial data prompted us to study whether m^6^A could serve as a mark that defines the functions of the HIV-1 full-length RNA as template for translation or packaging during viral replication.

Here, we show that methylation of two adenosine residues within the 5′-UTR by the METTL3/METLL14 complex inhibits full-length RNA packaging. m^6^A-seq analysis revealed that the full-length RNA present in purified viral particles lacks m^6^A at the 5′-UTR suggesting the existence of two populations that differ in their m^6^A patterns. Further bioinformatic analyses identified two highly conserved nucleotides A_198_ and A_242_ within the 5′-UTR as the key residues involved in the m^6^A-mediated regulation of gRNA packaging. We also observed that the full-length RNA is a substrate for the RNA demethylase FTO, which together with Gag drives RNA demethylation to promote packaging. Finally, the pharmacological targeting of FTO activity resulted in impaired full-length RNA metabolism and a strong inhibition of packaging. Together, our data reveal a novel mechanism by which Gag selects the molecules of gRNA that will be used for packaging, which is regulated by an epitranscriptomic switch within the 5′-UTR.

## RESULTS

### The presence of m^6^A alters the *in vitro* folding and dimerization of the HIV-1 full-length RNA 5′-UTR

Since previous studies suggesting that a conformational switch of the 5′-UTR could regulate the ability of the HIV-1 full-length RNA to function as mRNA or gRNA have not considered the presence of RNA modifications such as m^6^A, we first sought to determine the impact of adenosine methylation on the folding of the 5′-UTR. As a first approach to study the impact of m^6^A on RNA structure, we generated an *in* vitro-transcribed 5′-UTR containing unmodified adenosines or a 5′-UTR in which all adenosines were replaced by m^6^A. Both m^6^A-5′-UTR and A-5′-UTR were submitted to 1M7 SHAPE analysis in parallel as described in Methods. Reactivity towards SHAPE reagents reveals the ribose flexibility, and as a consequence, the pairing status of each nucleotide. As such, higher reactivity of a given nucleotide means a higher probability to be in a single strand conformation. Comparative analysis of the SHAPE reactivity profiles indicates that the presence of m^6^A significantly alters the folding of the 5′-UTR (Fig. 1a and Supplementary Fig. 1a). The first interesting observation from our SHAPE data is that we do not only observe a reactivity modification for As or Us.

**Figure 1:**
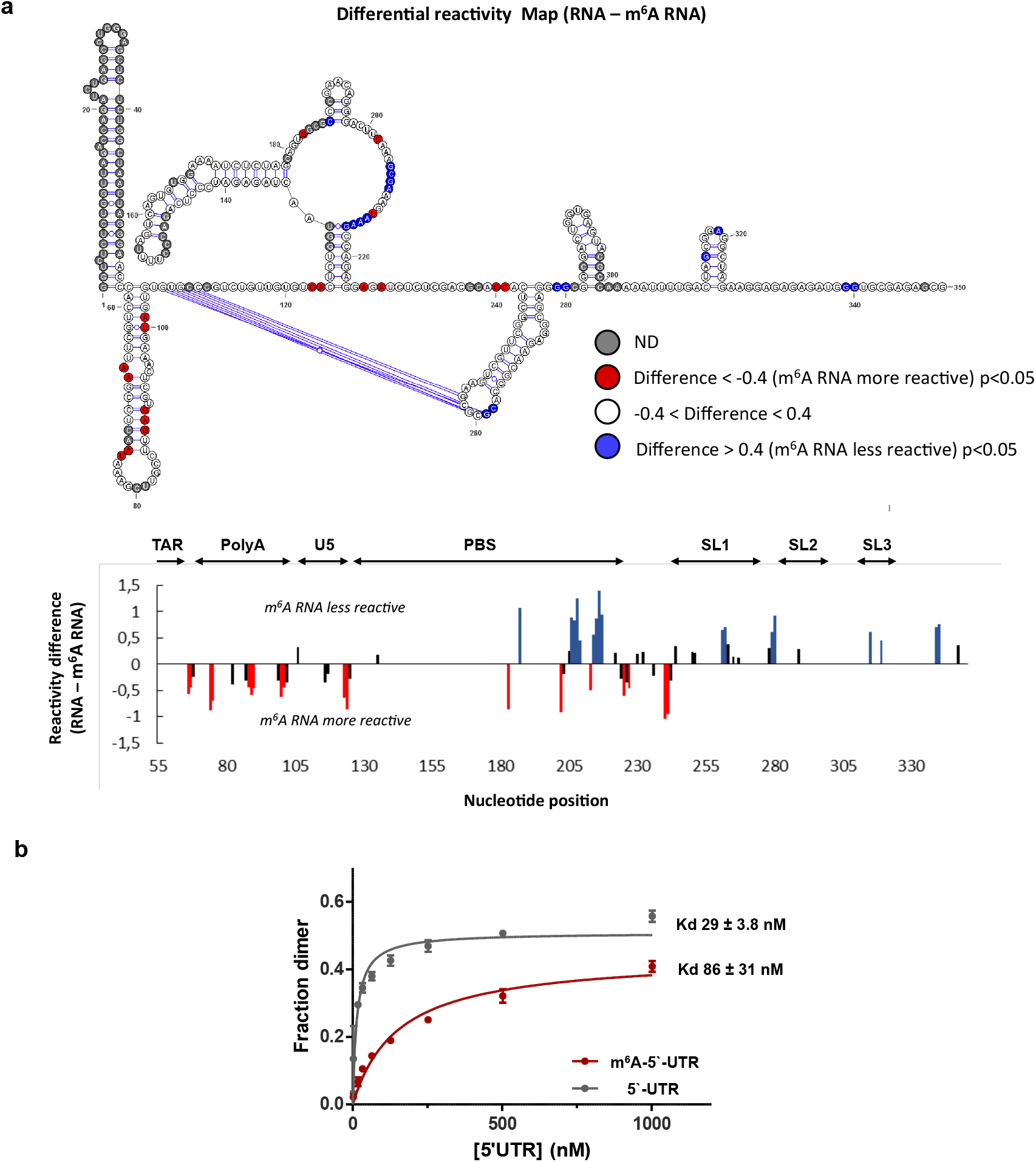
The presence of m^6^A alters the *in vitro* folding and dimerization of the HIV-1 full-length RNA 5′-UTR. **a,** SHAPE reactivity differences between *in vitro* transcribed 5′-UTR RNA containing 0% m^6^A or 100% m^6^A plotted on the secondary structure model of the HIV-1 full-length RNA 5′-UTR (upper panel). Nucleotides in red are significantly more reactive with m^6^A whereas those in blues are significantly less reactive (p-value < 0.05 and reactivity difference > 0.4). Nucleotides in white are of equivalent reactivity. Sites where the reactivity could not be determined are depicted in grey. The histogram shows the reactivity difference between 0% m^6^A 5′-UTR and 100% m^6^A 5′-UTR (lower). Blue and red bars highlight nucleotides with a significantly lower or higher reactivity in the 100% m6A RNA than in the 0% m6A RNA (p-value < 0.05 and reactivity difference > 0.4), respectively. Significative differences that are below the threshold of 0.4 are indicated in black. **b,** Fraction of dimers determined by electromobility shift assay for different concentrations of *in* vitro transcribed 5′-UTR (1 μM, 0.5 μM, 0.25 μM, 127 nM, 65 nM, 33 nM, 18 nM, 2 nM) harboring either 0% of m^6^A (grey) or 100%) of m^6^A (red) with the dissociation constants obtained from the data points (see Methods for details).

Moreover, although all adenosines were methylated in the m^6^A-5′-UTR RNA only local reactivity changes, often clustered, were observed suggesting that the presence of m^6^A influences the folding of specific domains or motifs within the 5′-UTR. The presence of m^6^A is predicted and has been observed to destabilize A-U pairings embedded in helical regions. This is mostly in agreement with the destabilizations (increase in reactivity) that are often observed for As, Us or in nucleotides in close proximity with proposed A-U base pairs. In particular, this could explain the reactivity enhancement within the poly-A stem, which is likely to be globally destabilized by m^6^A. However, none of the other nucleotides for which we observe a reactivity increase is involved or are close to A-U pairings and most of them are not even predicted to be base paired in most of the published models. In contrast, m^6^A has been shown to stabilize A-U pairings when preceded in 5′ by a bulged nucleotide. This does not offer a rationale for any of the reactivity drop we observe which are mostly predicted to be in single strand regions thus, suggesting that m^6^A modulates a higher order structure and/or tertiary pairings yet to be identified. This prompted us to monitor dimerization of the 5′-UTR and m^6^A-5′-UTR *in vitro*. We observed that the presence of m^6^A reduces but not abolish the efficiency of dimerization (Fig. 1b). The structure of the HIV-1 full-length RNA dimer is still poorly defined and a matter of debate, thus the dimerization deficiency observed does not provide a straightforward explanation for all the alterations of the SHAPE profile but could reveal unknown rearrangements.

Together, these data indicate that the presence of m^6^A may play an important role in the folding and dimerization of the 5′-UTR and prompted us to study this feature in a cellular context.

### The presence of m^6^A within the full-length RNA favors Gag synthesis but interferes with packaging

In order to study the role of m^6^A on the cytoplasmic fate of the full-length RNA during viral replication, we first determined the effects of METTL3/14 overexpression on Gag and full-length RNA levels obtained from cell extracts and purified viral particles (see scheme in Supplementary Fig. 2a). m^6^A-RIP analysis from METTL3/14 overexpressing cells showed an increase in the m^6^A/A ratio of the full-length RNA compared to the control indicating that the viral transcript is hypermethylated under these experimental conditions (Supplementary Fig. 2b). Consistent with the positive role of m^6^A on Gag synthesis previously described^20–22^, we observed that increased methylation of the full-length RNA by METTL3/14 overexpression results in increased levels of Gag and its processing products with minor effects on the intracellular levels of the full-length RNA (Fig. 2a). Quantification of viral particles produced from the same cells revealed a slight increase in Gag levels (as judged by anti-CAp24 ELISA) from METTL3/14 overexpressing cells, which could be attributed to the increased Gag synthesis observed (Fig. 2b).

**Figure 2:**
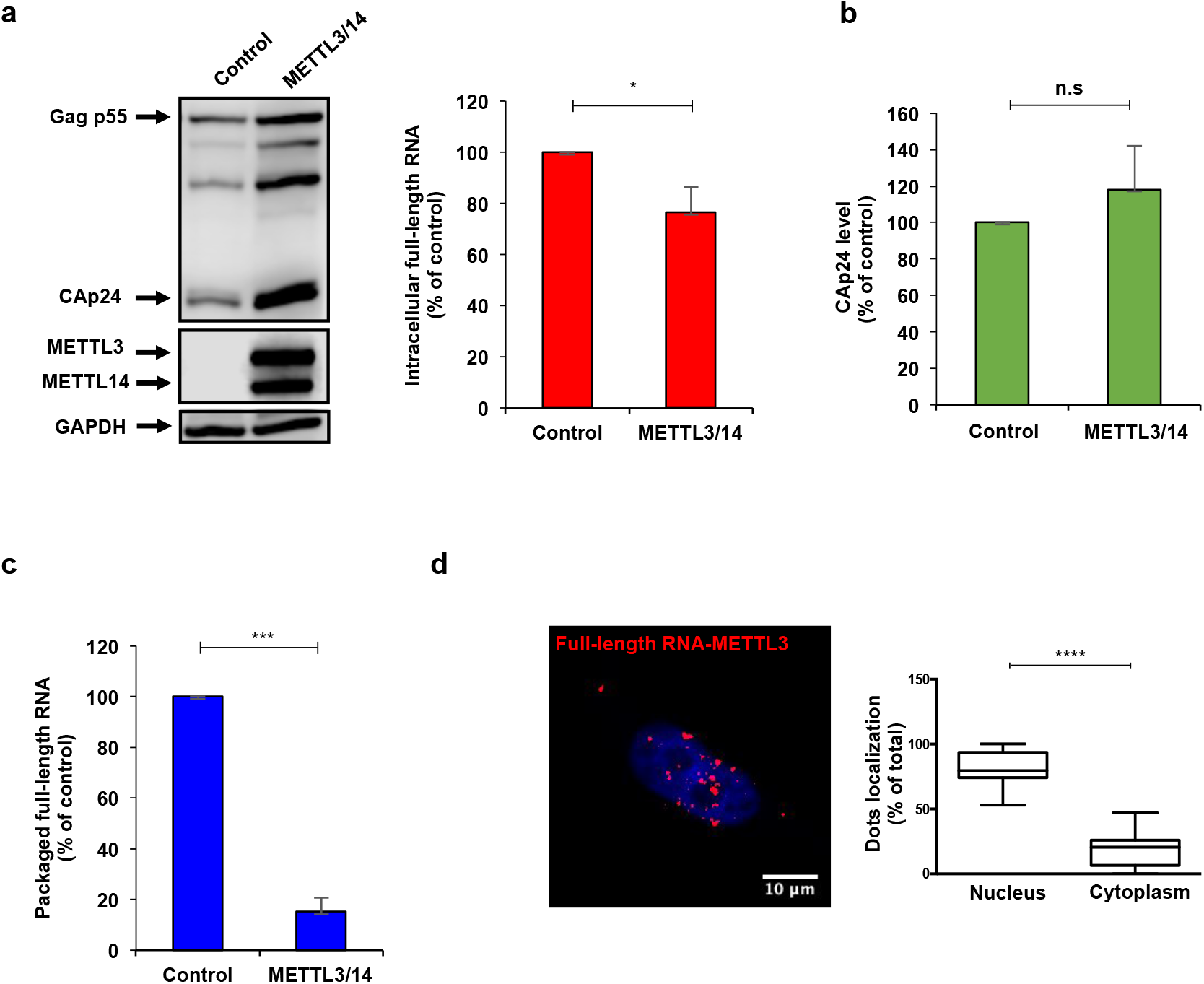
The presence of m^6^A within the full-length RNA favors Gag synthesis but interferes with packaging. HEK293T cells were transfected with pNL4.3 and pCMV-VSVg together with pCDNA-Flag-METTL3 and pCDNA-Flag-METTL14 or pCDNA-d2EGFP as a control, **a,** At 24 hpt cells extracts were used to detect Gag, Flag-METTL3 and Flag-METTL14 by Western blot. GAPDH was used as a loading control (left panel). In parallel, cells extracts were used to perform RNA extraction and the full-length RNA was quantified by RT-qPCR (right panel). Intracellular full-length RNA was normalized to the control (arbitrary set to 100%) and presented as the mean +/− SD of three independent experiments (\**P<0.05, t*-test). **b,** Supernatants from cell cultures in (a) were filtered and viral particles were purified by ultracentrifugation. The level of CAp24 was quantified by an anti-CAp24 ELISA. The level of CAp24 was normalized to the control (arbitrary set to 100%) and presented as the mean +/− SD of three independent experiments (n.s; non-significant, *t-* test). **c,** Viral particles purified from (b) were used to perform RNA extraction and the packaged full-length RNA from CAp24 equivalents was quantified by RT-qPCR. Packaged full-length RNA was normalized to the control (arbitrary set to 100%) and presented as the mean +/− SD of three independent experiments (\*\*\**P<0.001, t*-test). **d,** HeLa cells were transfected with pNL4.3, pCMV-VSVg and pCDNA-Flag-METTL3. At 24 hpt, the interaction between the full-length RNA and the Flag-tagged METTL3 was analyzed by ISH-PLA as described in the Methods section. Red dots indicate the interactions between the full-length RNA and the METTL3. Scale bar 10 mm. A quantification of the red dots in the nucleus (co-localizing with the DAPI staining) and the cytoplasm of 14 cells is presented on the right (\*\*\*\**P<0.0001*, Mann-Whitney test).

Then, we quantified the full-length RNA from equal amounts of viral particles and observed that viral particles produced under METTL3/14 overexpression contain around 3-fold less packaged gRNA indicating that hypermethylation of the full-length RNA impedes its packaging into nascent particles (Fig. 2c). We were not able to observe a similar effect of murine METTL3/14 overexpression on the simple retrovirus MLV, which was shown two segregate their full-length RNA into two specialized populations for translation and packaging further suggesting that MLV and HIV-1 might evolved diverse mechanisms for the cytoplasmic sorting of their full-length RNA (Supplementary Fig. 2c).

Since results presented above indicate that m^6^A deposition by METTL3/14 affects the cytoplasmic fate of the full-length RNA, we wanted to investigate where within the cell the m^6^A writer complex modifies the viral RNA. For this, we analyzed the interaction between the full-length RNA and METTL3 by *in situ* hybridization coupled to the proximity ligation assay (ISH-PLA)^24^. Confocal microscopy analyses revealed a predominant interaction within the nucleus, which suggests that the full-length RNA must be methylated in the nucleus and reach the cytoplasm in a methylated form (Fig. 2d and Supplementary Fig. 2d).

Together, these data suggest that nuclear methylation of the HIV-1 full-length RNA by METTL3/14 favors its use as mRNA for Gag synthesis but interferes with its incorporation into viral particles.

### Methylation of A_198_ and A_242_ within the 5′-UTR interferes with HIV-1 full-length RNA packaging

From data presented above, it seems that the presence of m^6^A interferes with the function of the HIV-1 full-length RNA as gRNA. Thus, to gain further insights into this regulation, we employed the m^6^A-seq strategy to determine the m^6^A patterns of the intracellular and viral particle-associated HIV-1 full-length RNA. In agreement with previous data reported for the NL4.3 and LAI.2 strains in T-lymphocytes and HEK293T cells^20–22^, we identified m^6^A peaks mainly at the 5′-UTR and a cluster of peaks at the 3′ end of the intracellular full-length RNA (Fig. 3a, see intracellular full-length RNA). Interestingly, we observed that the full-length RNA from purified viral particles maintains the m^6^A peak at the 3′-UTR but lacks the m^6^A peak at the 5′-UTR. This observation together with data from Fig. 2c suggests that the presence of m^6^A at the 5′-UTR interferes with the incorporation of the full-length RNA into viral particles (Fig. 3a, see packaged full-length RNA). Of note, this difference in the methylation patterns between intracellular and packaged RNA was not observed in the host 7SL RNA, which is also packaged at high levels into HIV-1 particles^25^, indicating a very specific effect of m^6^A on full-length RNA packaging (Supplementary Fig. 3a). These data strongly suggest that full-length RNA molecules lacking m^6^A at the 5′-UTR are primarily selected by Gag as gRNA to be incorporated into viral particles.

**Figure 3:**
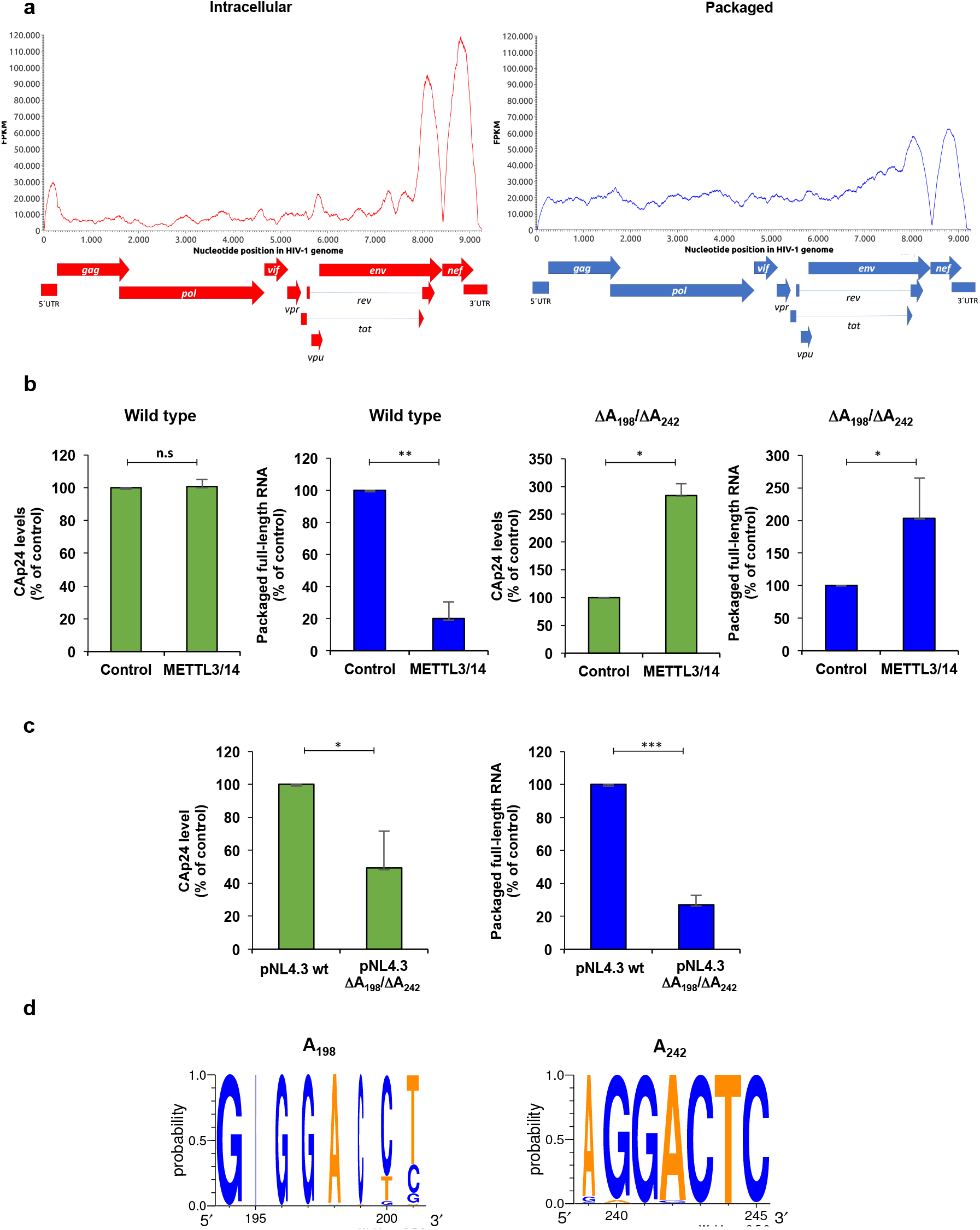
Methylation of A_198_ and A_242_ within the 5′-UTR interferes with HIV-1 full-length RNA packaging. **a,** HEK293T cells were transfected with pNL4.3 and pCMV-VSVg. Intracellular polyA RNA or viral particle-associated RNA was extracted at 24 hpt, fragmented and used for m^6^A-seq as described in Methods. Peak calling results for the intracellular (left) and packaged (right) full-length RNA is shown. **b,** HEK293T cells were transfected with pNL4.3 wild type or pNL4.3 DA_198_/DA_242_ together with pCMV-VSVg, pCDNA-Flag-METTL3 and pCDNA-Flag-METTL14 or pCDNA-d2EGFP as a control. At 24 hpt supernatants were filtered and viral particles were purified by ultracentrifugation. Purified viral particles were used to perform an anti-CAp24 ELISA and RNA extraction and RT-qPCR analysis as described above. The levels of CAp24 and the packaged full-length RNA (per CAp24 equivalents) were normalized to the control (arbitrary set to 100%) and presented as the mean +/− SD of three independent experiments (\**P<0.05*; \*\**P<0,01*; n.s, non-significant, *t*-test). **c,** HEK293T cells were transfected with pNL4.3 wild type or pNL4.3 DA_198_/DA_24_2 together with pCMV-VSVg and the supernatant was filtered and ultracentrifuged at 24 hpt. Purified viral particles were used to perform an anti-CAp24 ELISA and for RNA extraction and RT-qPCR analysis as described above. The levels of CAp24 and the packaged full-length RNA (per CAp24 equivalents) were normalized to the control (arbitrary set to 100%) and presented as the mean +/− SD of three independent experiments (\**P<0.05*; \*\*\**P<0,001, t*-test). **d,** Conservation analyses of adenosines 198 and 242 from 879 sequences from the HIV-1 database.

A bioinformatic prediction of the potentially methylated residues within the 5-UTR of the NL4.3 strain identified A_198_ and A_242_ in a very favorable methylation context (Supplementary Fig. 3b). Both residues are contained within the m^6^A peak we have identified within the 5′-UTR of the intracellular full-length RNA (Supplementary Fig. 3c) and were also identified in previous m^6^A-seq data obtained from T-lymphocytes and HEK293T cells^20, 22^. Thus, we deleted A_198_, A_242_ or both from the NL4.3 provirus in order to determine the role of these adenosine residues on the m^6^A-mediated regulation of full-length RNA packaging. We observed that ΔA_198_ and ΔA_242_ single mutant proviruses were slightly resistant to the effects of METTL3/14 overexpression on full-length RNA packaging (Supplementary Fig. 3d). However, the ΔA_198_/ΔA_242_ double mutant provirus was refractory to the positive effects of METTL3/14 overexpression on Gag synthesis and the negative effects on full-length RNA packaging observed with the wild type provirus indicating that A_198_ and A_242_ are key residues involved in this m^6^A-mediated regulation (Fig. 3b and Supplementary Fig. 3e). Next, we sought to investigate the role of A_198_ and A_242_ on HIV-1 full-length RNA metabolism. A comparison between wild type and the ΔA_198_/A_242_ provirus showed that the double mutant virus accumulates more intracellular Gag but releases significantly less viral particles (Fig. 3c and Supplementary Fig. 3f). Moreover, quantification of the full-length RNA from equal amounts of viral particles revealed a defect in packaging in the ΔA_198_/A_242_ double mutant provirus thus, confirming the critical relevance of these two adenosine residues for the incorporation of the HIV-1 full-length RNA into viral particles (Fig. 3c). An analysis of 890 sequences from the HIV database (www.hiv.lanl.gov) indicate that A_198_ and A_242_ are highly conserved within the 5′-UTR of isolates suggesting that this epitranscriptomic regulation must be a common feature of different HIV-1 subtypes including the highly prevalent subtypes C and B as well as circulating recombinant forms (Fig. 3d and Supplementary Fig. 3g). Taking together, these results indicate that the HIV-1 full-length RNA may exist as two different populations that differ at least in the m^6^A residues present within the 5′-UTR. Only the full-length RNA molecules lacking m^6^A at positions 198 and 242 might be recognized by Gag for packaging.

### Demethylation by a Gag-FTO complex favors HIV-1 full-length RNA packaging

Then, we sought to determine whether this m^6^A-mediated regulation of full-length RNA packaging was a dynamic process. This was important considering that the reversible nature of adenosine methylation in cellular mRNA has been challenged^26^. For this, we overexpressed the RNA demethylase FTO and the analysis of the m^6^A/A ratio of the full-length RNA in control and FTO overexpressing cells revealed that the viral RNA is indeed a substrate for this m^6^A eraser (Supplementary Fig. 4a). Consistent with a positive role of m^6^A on Gag synthesis, we observed that FTO-induced demethylation of the full-length RNA results in a reduction of Gag levels despite a slight increase in intracellular full-length RNA levels (Fig. 4a). We also observed minimal changes in the CAp24 levels from purified viral particles produced under RNA demethylation conditions (Fig. 4b). However and in agreement with a negative role of m^6^A on full-length RNA packaging, we observed that viral particles produced from FTO overexpressing cells contain around 3-fold more packaged gRNA compared to the control (Fig. 4c). It should be mentioned that we were not able to observe similar results with the RNA demethylase ALKBH5 suggesting that full-length RNA demethylation by FTO is important for packaging (Supplementary Fig. 4b).

**Figure 4:**
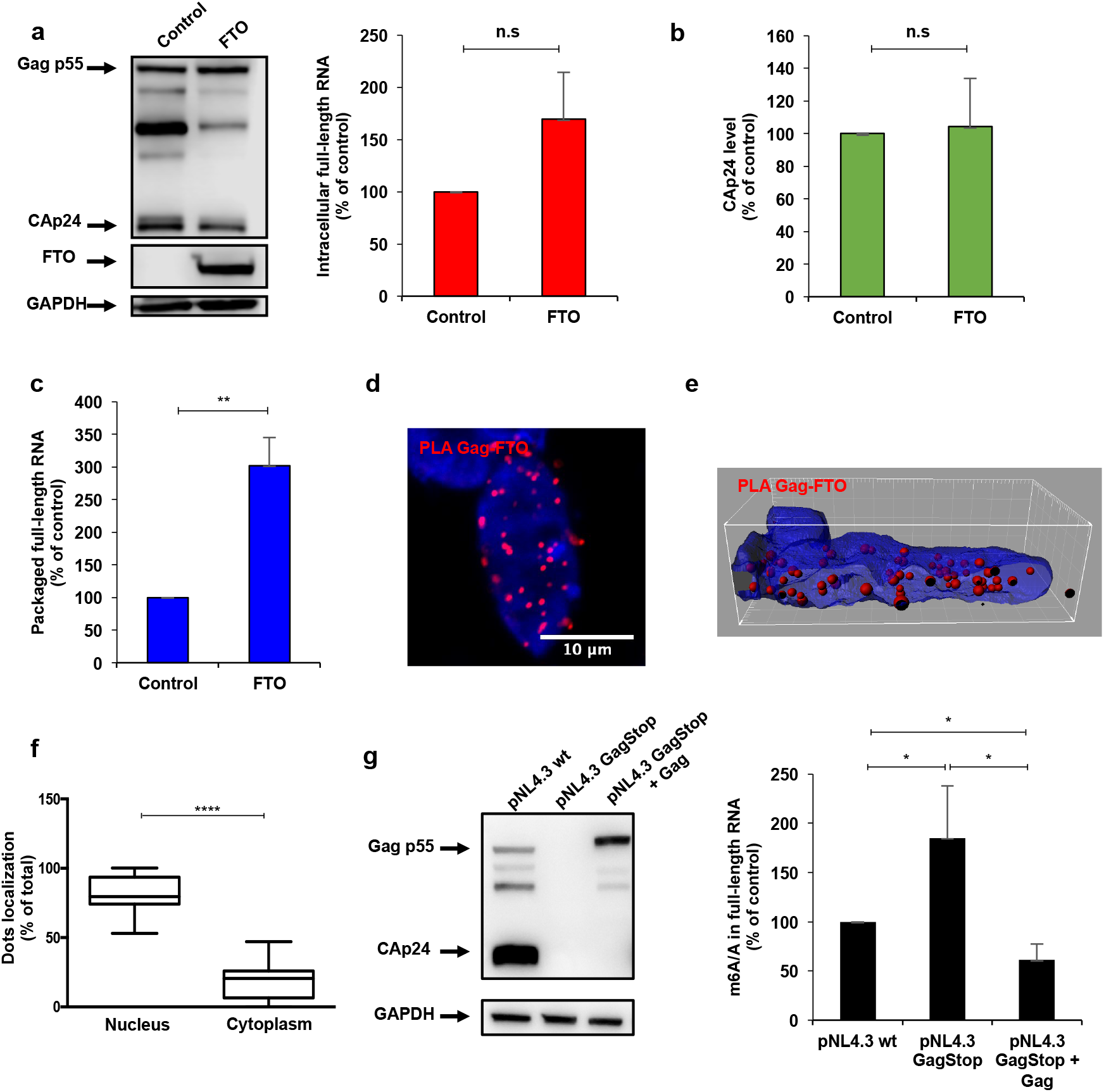
Demethylation by a Gag-FTO complex favors HIV-1 full-length RNA packaging. HEK293T cells were transfected with pNL4.3 and pCMV-VSVg together with pCDNA-3XFlag-FTO or pCDNA-3XFlag-d2EGFP as a control. **a,** At 24 hpt cells extracts were used to detect Gag and 3XFlag-FTO by Western blot. GAPDH was used as a loading control (left panel). In parallel, cells extracts were used to perform RNA extraction and the full-length RNA was quantified by RT-qPCR (right panel). Intracellular full-length RNA was normalized to the control (arbitrary set to 100%) and presented as the mean +/− SD of three independent experiments (\**P<0.05, t*-test). **b,** Supernatants from cell cultures in (a) were filtered and viral particles were purified by ultracentrifugation. The level of CAp24 was quantified by an anti-CAp24 ELISA, normalized to the control (arbitrary set to 100%) and presented as the mean +/− SD of three independent experiments (n.s; non-significant, *t*-test). **c,** Viral particles purified from (b) were used to perform RNA extraction and the packaged full-length RNA from CAp24 equivalents was quantified by RT-qPCR. Packaged full-length RNA was normalized to the control (arbitrary set to 100%) and presented as the mean +/− SD of three independent experiments (\*\**P<0.01, t*-test). **d,** HeLa cells were co-transfected with pNL4.3, pCMV-VSVg and pCDNA-3XFlag-FTO. At 24 hpt, the interaction between Gag and Flag-tagged FTO was analyzed by PLA as described in Methods. Red dots indicate the interactions between Gag and FTO (left panel). Scale bar 10 mm. **e,** Three-dimensional reconstitution of the PLA results shown in (d) was performed to determine the subcellular localization of the interaction between Gag and FTO. **f,** Quantification of the red dots in the nucleus (co-localizing with the DAPI staining) and the cytoplasm of 15 cells is presented on the right (\*\*\*\**P<0,0001*, Mann-Whitney test). **g,** HEK293T cells were transfected with the pNL4.3 wild type, pNL4.3-GagStop or pNL4.3-GagStop together with pCDNA-Gag. At 24 hpt cells extracts were used to detect Gag and GAPDH was used as a loading control. In parallel, cells extracts were used to perform RNA extraction followed by an immunoprecipitation using an anti-m^6^A antibody (m^6^A-RIP as described in Methods). The full-length RNA from the input (“A” fraction) and from the immunoprecipitated material (“m^6^A” fraction) was quantified by RT-qPCR. The m^6^A/A ratio was normalized to pNL4.3 wild type (arbitrary set to 100%) and presented as the mean +/− SD of three independent experiments (\**P<0.05, t*-test).

Considering that m^6^A demethylation favors full-length RNA packaging, we wanted to know where within the cell the viral RNA became demethylated by FTO. For this, we analyzed the interaction of the between the full-length RNA and FTO in cells by ISH-PLA but despite several attempts, we were not able to detect a direct interaction regardless all the components were correctly expressed within the cells (Supplementary Fig. 4c). This observation suggests that either there is no a massive interaction between the full-length RNA and FTO or that such interactions occurs very transiently (or at very low rates) being below the detection limit of our ISH-PLA strategy.

The lack of a detectable interaction between the full-length RNA and FTO prompted us to investigate whether Gag could interact with FTO and drive full-length RNA demethylation. In order to test this possibility, we employed the proximity ligation assay (PLA) and observed that Gag and FTO indeed form complexes in cells (Fig. 4d). Interestingly, quantification of the dots per cell localizing with the nuclear staining as well as 3D reconstitutions of representative images indicate that Gag and FTO mostly associates within the nucleus (Figs. 4e and 4f). Indeed, we observed that FTO overexpression increases the nuclear localization of Gag (Supplementary Fig. 4d).

From these results, it was tempting to speculate that the full-length RNA is methylated within the nucleus by METLL3/14 and Gag interacts with FTO in the nucleus to drive demethylation of the full-length RNA molecules that will be incorporated into assembling viral particles. Thus, we analyzed the m^6^A content of the full-length RNA in the presence or absence of Gag by using the wild type NL4.3 provirus and a mutant provirus containing premature stops codons that abolish Gag synthesis (GagStop provirus). Compared to the wild type full-length RNA, the level of m^6^A increases when Gag is absent and is restored or even decreased when the Gag protein was expressed *in trans* indicating that Gag regulates the methylation status of the full-length RNA (Fig. 4g).

Together, these results strongly indicate that the FTO-mediated demethylation is required for full-length RNA packaging in a process supported by the Gag precursor.

### Inhibition of FTO demethylase activity impacts full-length RNA metabolism and blocks packaging

We finally sought to determine whether this epitranscriptomic regulation of the HIV-1 full-length RNA packaging was a potential therapeutic target for pharmacological intervention. For this, we took advantage of the ester form of meclofenamic acid (MA2), which was shown to specifically interfere with FTO-mediated m^6^A demethylation^27^. Therefore, we analyzed Gag and the full-length RNA in cells treated with DMSO (as a control) or MA2 and observed a reduction in Gag synthesis and the intracellular levels of the full-length RNA indicating that FTO-mediated demethylation is required for proper metabolism of the full-length RNA within the cell (Fig. 5a). Consistent with a perturbed intracellular full-length RNA metabolism, we observed a decrease in the viral particles released from MA2-treated cells (Fig. 5b). Strikingly, quantification of the packaged full-length RNA from equal amounts of viral particles indicates that inhibition of FTO activity by MA2 almost abolished packaging (Fig. 5c).

**Figure 5:**
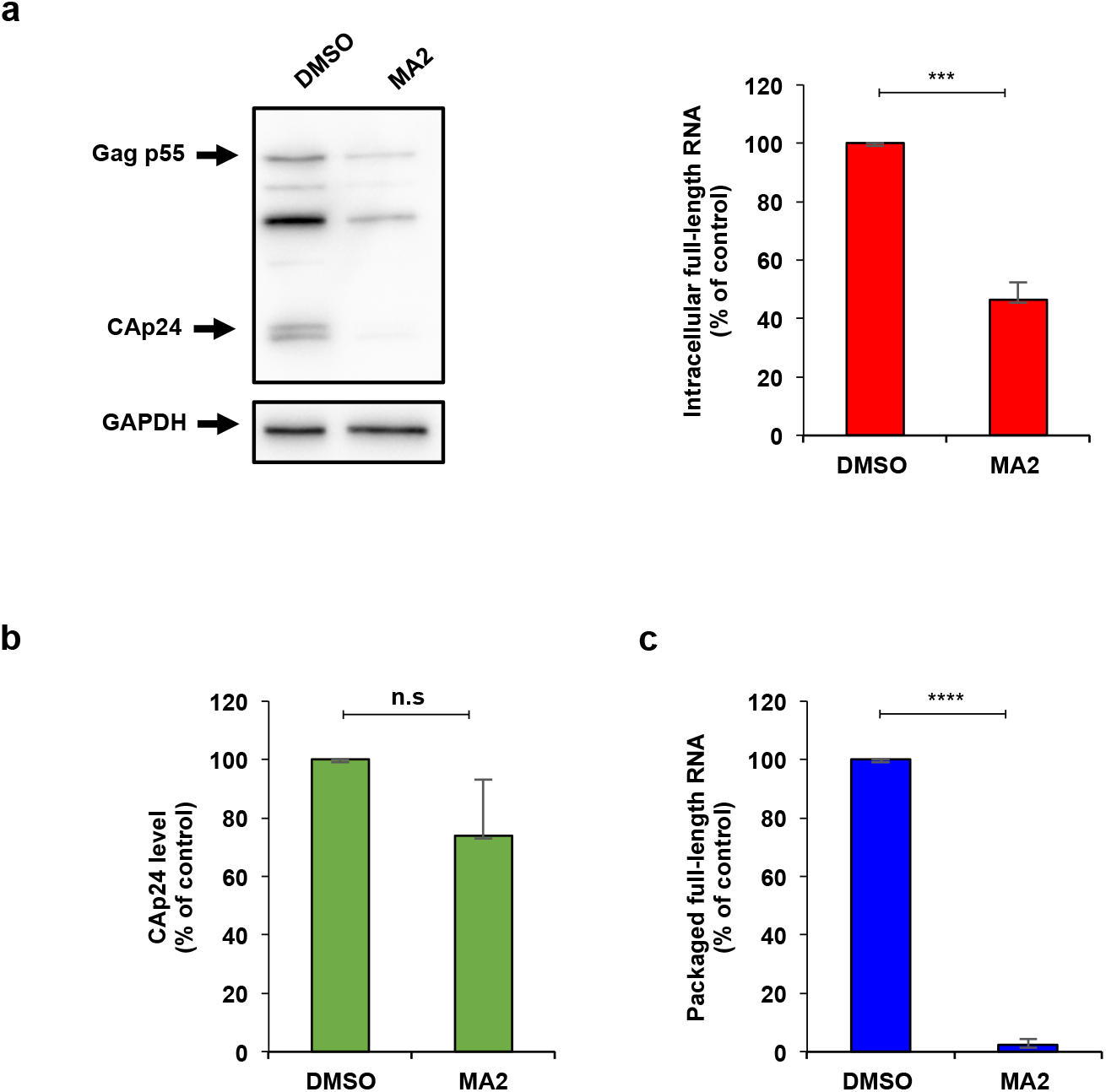
Inhibition of FTO demethylase activity impacts full-length RNA metabolism and blocks packaging. HEK293T cells were transfected with pNL4.3 and pCMV-VSVg and were treated with MA2 or DMSO as a control. **a,** At 24 hpt cells extracts were used to detect Gag and GAPDH as a loading control (left panel). In parallel, cells extracts were used to perform RNA extraction and the full-length RNA was quantified by RT-qPCR (right panel). The intracellular full-length RNA was normalized to the control (arbitrary set to 100%) and presented as the mean +/− SD of three independent experiments (\*\*\**P<0,001, t*-test). **b,** At 24 hpt the supernatant was filtered and viral particles were purified by ultracentrifugation. The level of CAp24 was quantified by an anti-CAp24 ELISA, normalized to the control (arbitrary set to 100%) and presented as the mean +/− SD of three independent experiments (n.s; non-significant, *t*-test). **c,** Purified viral particles from (b) were used to perform an RNA extraction and the packaged full-length RNA from CAp24 equivalents was quantified by RT-qPCR. Packaged full-length RNA was normalized to the control (arbitrary set to 100%) and presented as the mean +/− SD of three independent experiments (\*\*\*\**P<0.0001, t*-test).

Taking together, these results confirm that FTO-mediated demethylation is critical for HIV-1 full-length RNA packaging and this process is a potential target for the design of novel antiretroviral drugs.

## DISCUSSION

Assembly of human immunodeficiency virus type-1 particles is a highly regulated process in which the major structural polyprotein Gag together with other viral and cellular components are recruited to the plasma membrane for the release of the viral progeny. The assembly process occurs in multiple steps driven by the different functional domains that compose the Gag precursor. As such, while the nucleopcapsid (NC) domain specifically recruits two copies of the full-length RNA, the matrix (MA) domain allows targeting of the complex to specific plasma membrane micro-domains and the capsid (CA) domain drives Gag multimerization at such sites. Packaging of two copies of full-length RNA by the NC domain of Gag is highly specific and occurs selectively over thousands of cellular and viral RNA species. This selectivity was proposed to be possible by the presence of cis-acting RNA signatures spanning the 5′-UTR and the beginning of the Gag coding region. However, the full-length RNA also serves as mRNA for the synthesis of Gag and Gag-Pol precursors and thus, translation and packaging are expected to be two mutually exclusive processes. Although the adoption of a branched multiple hairpin (BMH) conformation of the 5′-UTR was initially proposed to favor dimerization and packaging over translation^12–14^, it was later demonstrated that translation of the full-length RNA is under positive selection and thus, not regulated by a conformational switch of the 5′-UTR^18, 19^. Additional structural studies carried out in cells and virions also argued against structural rearrangements as drivers of the transition between translation and packaging of the HIV-1 full-length RNA^16, 17, 28^. Therefore, the mechanism by which Gag selects “packageable” from “translatable” full-length RNA molecules still remains as one of the long-lasting questions in Retrovirology. In this work, we showed that demethylation of two highly conserved adenosine residues within the 5′-UTR is critical for packaging of the HIV-1 full-length RNA. Interestingly, we observed that Gag associates with the RNA demethylase FTO in the nucleus and promotes demethylation of the full-length RNA, suggesting that Gag may drive FTO-mediated demethylation of those RNA molecules that will be used as gRNA to be incorporated into assembling viral particles (Fig. 6). This differential epitranscriptomic regulation exerted on the full-length RNA depending on its functions (mRNA or gRNA) may also help to explain the controversies reported in the literature^29^. As such, while the presence of m^6^A favors Gag synthesis through YTHDF proteins acting on the full-length RNA molecules destined to serve as mRNA^21^, the same cytoplasmic m^6^A readers may recognize specific features and drive degradation of the incoming viral RNA early upon infection (i.e., when the full-length RNA acts as gRNA)^22^.

**Figure 6.**
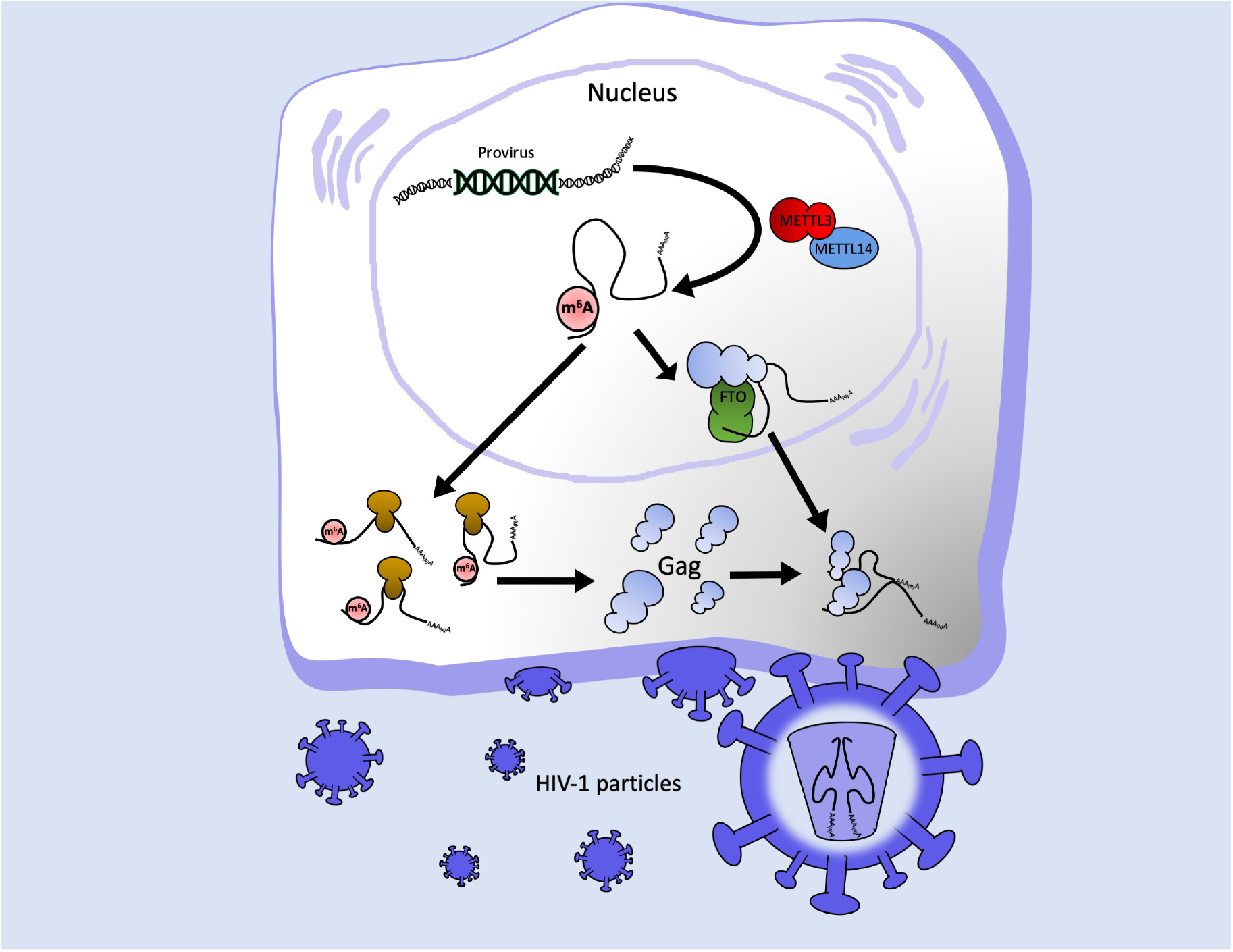
Working model for the epitranscriptomic regulation of HIV-1 full-length RNA packaging. The HIV-1 full-length RNA is methylated by METTL3/14 complex in the nucleus (for simplicity, only the presence of m^6^A on the 5′-UTR is shown). However, the structural protein Gag interacts with the m^6^A eraser FTO and drives demethylation of adenosines residues present at the 5′-UTR in a process required for full-length RNA packaging.

Further studies are required to elucidate the mechanism by which the binding of YTHDF proteins to m^6^A residues negatively impact full-length RNA metabolism in the absence of translation and whether methylation of the 5′-UTR is involved. In addition, the molecular mechanism by which the presence of m^6^A at the 5′-UTR interferes with full-length RNA packaging also deserves further investigation. One of the most plausible explanations is that recognition of m^6^A residues at the 5′-UTR by reader proteins interferes with Gag binding and/or full-length RNA dimerization. However, it is also possible that the presence of m^6^A itself directly repeals the recruitment of Gag or alters the optimal RNA conformation of the dimerization and/or packaging signal. Indeed, our *in vitro* structural and dimerization analyses support this last idea as they suggest that the presence of m^6^A affects the folding and dimerization of the 5′-UTR. Of note, although neither A_198_ nor A_242_ showed a significant reactivity alteration this does not exclude that their modification may have influenced the reactivity of close nucleotides such as G_240_ and G_241_. To date no studies have monitored the effect of m^6^A on such a complex structure and here we clearly show that this modification can have not only local but more global effects on RNA folding. Indeed, we observed that the introduction of m^6^A alters dimerization of the 5′-UTR, which could at least partly explain why methylated full-length RNA are not recovered within viral particles. Nevertheless, our experiments using the ΔA_198_/ΔA_242_ provirus strongly indicate that these two adenosine residues play a major role in packaging. While A_198_ is located within the region complementary to the tRNA^Lys3^ at the primer binding site (PBS), A_242_ is located in the AGGA bulge at the base of the SL1 region and corresponds to a Gag-binding domain previously shown by chemical probing *in vitro* and *in virio* as well as by CLIP-seq studies^17, 30, 31^. Further studies are required to fully understand the role of these residues in full-length RNA packaging.

In contrast to the simple retrovirus MLV, which segregates its full-length RNA into two separate populations for translation and packaging, the full-length RNA of the human lentivirus HIV-1 was proposed to exist as a single population that can indistinctly serve as mRNA or gRNA. While the presence of two specialized full-length RNA populations supports the lack of an epitranscriptomic regulation of MLV packaging, our data obtained with HIV-1 suggest that its full-length RNA may also exist as two populations with different m^6^A patterns. From these two populations, those molecules lacking m^6^A at the 5′-UTR will be preferentially selected by Gag for packaging (Fig. 6). In this regard, the HCV core protein was also shown to bind preferentially to RNA molecules lacking m^6^A most probably to avoid YTHDF proteins-mediated degradation upon viral entry^32^. Whether the packaging of hypomethylated RNA genomes is a conserved mechanism evolved by RNA viruses to avoid early degradation by cytoplasmic m^6^A readers must be determined.

Although the reversibility of adenosine methylation as well as the main target of the RNA demethylase FTO have been recently challenged^26, 33^, we showed that the HIV-1 full-length RNA is a substrate for FTO and this RNA demethylase regulates the incorporation of the viral genome into released viral particles. Interestingly, treatment of HIV-1 producer cells with the specific FTO inhibitor meclofenamic acid resulted in an impairment of full-length RNA metabolism with a potent effect on packaging, confirming the critical role of the demethylase activity of FTO during viral replication. In addition to meclofenamic acid, two small molecules developed by structure-based rational design were recently described as specific inhibitors of FTO m^6^A demethylase activity with the potential to be used as a treatment for adult myeloid leukemia^34^. Therefore, this novel epitranscriptomic mechanism regulating packaging of the HIV-1 full-length RNA could also be exploited as a target for pharmacological intervention.

## METHODS

### Selective 2′-hydroxyl acylation analyzed by primer extension (SHAPE)

The HIV-1 full-length RNA 5′-UTR was *in vitro* transcribed using the T7 RNA polymerase as described^35^. For m^6^A RNAs, ATP was substituted by N^6^-methyl-ATP (Jena Bioscience). RNA was quantified by measurement of the OD_256_ using a BioSpec-nano (Shimadzu). RNA integrity was assessed by agarose gel electrophoresis. SHAPE probing was conducted essentially as in^17^ with minor modifications. Briefly, 6 pmol of *in vitro* transcribed RNA containing 100% m^6^A or 0% m^6^A were diluted in 24 μl of water, denatured at 80°C for 2 minutes and ice cooled. After addition of 3 μl of 10X Folding Buffer (HEPES Ph7.5 400 nM, KCl 1M, MgCl_2_ 50 mM), samples were incubated for 10 minutes at room temperature and then 10 minutes at 37°C. Then, the RNA solution was added to 3 μl of 20 mM 1M7 (2mM final) (AEchem Scientific Corporation) or to 3 μl of DMSO (control) and incubated for 6 minutes at 37°C. RNA was subsequently precipitated in presence of 1 μl of 20 mg/ml of glycogen, 3 μl of 5M sodium acetate, 100 μl of ETOH for 1 hour at −20°C and then washed with ETOH 70% and resuspended in water. For primer extension, RNA was treated with 1 μl of DMSO and denatured for 3 minutes at 95°C and cooled at 4°C. Samples were mixed with 3 μl of 2μM fluorescent primer (D4-5′-TTTCTTTCCCCCTGGCCTT for the probed/control samples and D2-5′-TTTCTTTCCCCCTGGCCTT for the sequencing reaction, Sigma) and incubated for 5 minutes at 65°C, 10 minutes at 35°C and then 1 minute on ice. 5 μl of Reverse Transcription Mix (10 μl of 10 mM dNTPs and 40 μl of 5X MMLV RT Buffer Promega), 1 μl of 10 mM ddTTP for the sequencing reaction and 1 μl of MMLV Reverse Transcriptase RNAse H minus (Promega) were finally added and the reverse transcription was performed at 35°C for 2 minutes, 42°C for 30 minutes and 55°C for 5 minutes. Samples were then ice cooled and precipitated with ETOH for 2 hours. Pellets were resuspended in 40 μl of Sample Loading Solution (Beckman Coulter). Reverse transcription products were resolved on a CEQ-8000 sequencer (Beckman Coulter). Electropherograms were analyzed using QuSHAPE^36^. Raw data were processed by excluding the 2% of the highest values and normalizing the remaining values by the mean of the next 8% highest values^37^. The experiments were performed three times and reproducibility was assessed by calculating the standard error of the mean. Secondary structure was drawn using VaRNA^38^.

### Dimerization assay

*In vitro* transcribed 5′-UTR was serially diluted in 10 mM Tris pH 7, 10 mM NaCl, 140 mM KCl to obtain a final concentration ranging from 0 to 1 μM. 20 fmol (2 nM final) of radiolabeled RNA was added. Samples were denatured at 95°C for 3 minutes and then ice cooled. After addition of 1 mM MgCl_2_, RNA was allowed to fold for 30 minutes at 37°C. Samples were subsequently chilled on ice, mixed with a 5X native loading buffer (glycerol 20%, xylene cyanol 0.1%, bromophenol blue 0.1%) and loaded on a native 4% acrylamide gel. Samples were run for 1 hour at 100V on 4% native acrylamide mini-gels containing 34 mM Tris, 54 mM HEPES, 0.1 mM EDTA and 2.5 mM MgCl_2_. Dried gels were quantified using a BAS-5000 phosphorimager and MultiGauge 3.0 (Fujifilm). The fraction of dimer was calculated as the ratio “dimer – background” over “dimer + monomer – 2*background”. Data were fitted to Fraction dimer = Bmax*[RNA]/(Kd+[RNA]) using Prism 5.02 (GraphPad Software).

### Cell culture, DNA transfection and viral particle purification

HEK293T and HeLa cells were maintained in DMEM (Life technologies) supplemented with 10% FBS (Hyclone) and antibiotics (Hyclone) at 37°C and 5% CO_2_ atmosphere. Cells growing in 6-well plates (2.5×10^5^ cells/well) were transfected using linear PEI ~25000 Da (Polyscience) as described previously^24^ Cells were transfected using a ratio μg DNA/μl PEI of 1/15 and the DNA/PEI mix was incubated for 20 min at room temperature before adding to the cells. For experiments involving the FTO inhibitor, the culture medium was replaced by medium containing dimethyl sulfoxide (DMSO) as a control or 80 μM of an ethyl ester form of meclofenamic acid diluted in DMSO (MA2) prior DNA transfection. For viral particle purification, the supernatant was collected and filtered by passing through a 0.22 μm filter and then ultracentrifugated at 25.000 rpm for 2 hours at 4°C in a 20% sucrose cushion (prepared previously and stored at 4 °C). Purified viral particles were resuspended in 100 μl of PBS and stored in aliquots at −80 °C to then perform anti-CAp24 ELISA (HIV-1) or Western blot (MLV) or RNA extraction. Cells were also collected to perform Western blot and RNA extraction as described in Supplementary Methods.

### m^6^A-seq

Poly(A) RNA was purified from 100 μg of total RNA extracted from HEK293T cells previously transfected with pNL4.3 and pCMV-VSVg. Briefly, total RNA in 500 μl of water was incubated at 65°C for 10 minutes and incubated with 3 μl of oligo dT-Biotin (dT-B; IDT Technologies) (50 pmol/μl) and 13 μl of SSC Buffer 20X (Santa Cruz Biotechnology) and allowed to cool at room temperature. 600 μg of Dynabeads™ Streptavidin (60 μl; Thermo Fisher) were washed three times with Buffer SSC 0.5X and resuspended in 100 μl of Buffer SSC 0.5X. Then, the RNA/oligo dT-B mix was incubated with the streptavidin beads at room temperature for 10 minutes in head-over-tail rotation. RNA-beads were washed four times with 300 μl Buffer SSC 0.1X and bound RNA was eluted twice with 100 μl of water. The RNA was precipitated with 10 mM MgCl_2_, 20 μg glycogen (Thermo Fisher) and 2.5 volume of ETOH 100% overnight at −20 °C and then washed with ETOH 70%. Poly(A) RNA as well as RNA obtained from purified viral particle were fragmented using Fragmentation Reagent (Thermo Fisher). For this, 2 μg of RNA in 9 μl of water was incubated with 1 μl of Fragmentation Buffer 10X for 15 minutes at 70 °C, then 1 μl of Stop solution was added and incubated on ice. The RNA fragmented was precipitated overnight as described above. Fragmented RNA diluted in 380 μl was heated at 70 °C for 5 minutes, placed on ice for 3 minutes. The denatured RNA was mixed with 1 μl of rRNasin^®^ (Promega), 5 μl VRC, 100 μl of IP Buffer 5X (50 mM Tris-Hcl pH7.4, 750 mM NaCl and 0.5% NP-40) and 5 μl of an anti-m^6^A antibody (0,5 mg/mL; Synaptic System #202003) and incubated for 2 hours at 4 °C with head-over-tail rotation. At the same time, 600 μg of Dynabeads™ Protein A magnetic beads (20 μl; Thermo Fisher) were washed in 1 ml of IP Buffer 1X with 1 μl of VRC and were incubated with 500 μl of Buffer IP 1X with 0.5 mg/ml of BSA for 2 hours at 4 °C with head-over-tail rotation. Then, beads were washed with 500 μl of IP Buffer 1X and added to the RNA/anti-m^6^A antibody mix. The RNA-beads mix was incubated for 2 hours at 4°C in head-over-tail rotation. After incubation, the RNA-beads mix was washed twice with 500 μl IP Buffer 1X. Bound RNA was eluted with 100 μl of Elution Buffer (5mM Tris-HCl, 1mM EDTA and 0.05% SDS) and 1 μl of Proteinase K (New England BioLabs) and incubated for 1.5 hour at 50°C. RNA was extracted from supernatant using TRIzol™ (ThermoFisher). The RNA recovered was precipitated with 10 mM MgCl_2_, 20 μg glycogen (Thermo Fisher) and 2.5 volumes of ETOH 100% overnight at −20 °C and then washed with ETOH 70%. Equal amounts of RNA from input and immunoprecipitation were used for RT-qPCR. cDNA libraries preparations and RNAseq was performed at Genoma Mayor. All the samples were sequenced in an Illumina HiSeq2000 platform with paired-end 100 bp read length. Read quality was evaluated with *Fastqc* and the *Burrows-Wheeler Alignment Tool (BWA-Mem)* was used for mapping reads to the HIV-1 genome with default parameters.

The alignment data were analyzed with *MACS2* to call peaks with -*f BAMPE --nomodel -- SPMR* options for generated viral peaks data and to generate FPKM (Fragments Per Kilobase per Million mapped reads). Predicted peaks were sorted by average coverage.

## DATA AVAILABILITY

m^6^A-seq data were deposited at the GEO upon accession number GSE130687.

## AKNOWLEDGEMENTS

Authors wish to thank Professor Cai-Guang Yang (CAS Key Laboratory of Receptor Research, Shangai Institute of Materia Medica, Chinese Academy of Sciences) for providing MA2. Authors will also thank to Dr. Chuan He (University of Chicago), Dr. Yun-Gui Yang (Beijing Institute of Genomics, Chinese Academy of Sciences), Dr. Jin Crystal Zhao and Dr. Monica Roth (University of Wisconsin) for providing expression vectors as well as Dr. Gloria Arriagada (Universidad Andrés Bello) for providing the anti-MLV CAp30 antibody.

## AUTHOR CONTRIBUITIONS

CP-M designed and performed experiments, analyzed data and wrote the paper. DT-A, SR-B, BR-A, FGdeG, PA-C, CA-S, MLA and JCh performed experiments and analyzed data. CR-F performed bioinformatic analyses. GdeB performed *in vitro* experiments and analyzed data. FV-E and BS contributed to data analysis and manuscript writing. RS-R contributed to the concept and design, data analysis and manuscript writing. All the authors read and approved the final version of the manuscript.

