## Supplementary Figures for "An epitranscriptomic switch at the 5′-UTR controls genome selection during HIV-1 genomic RNA packaging"

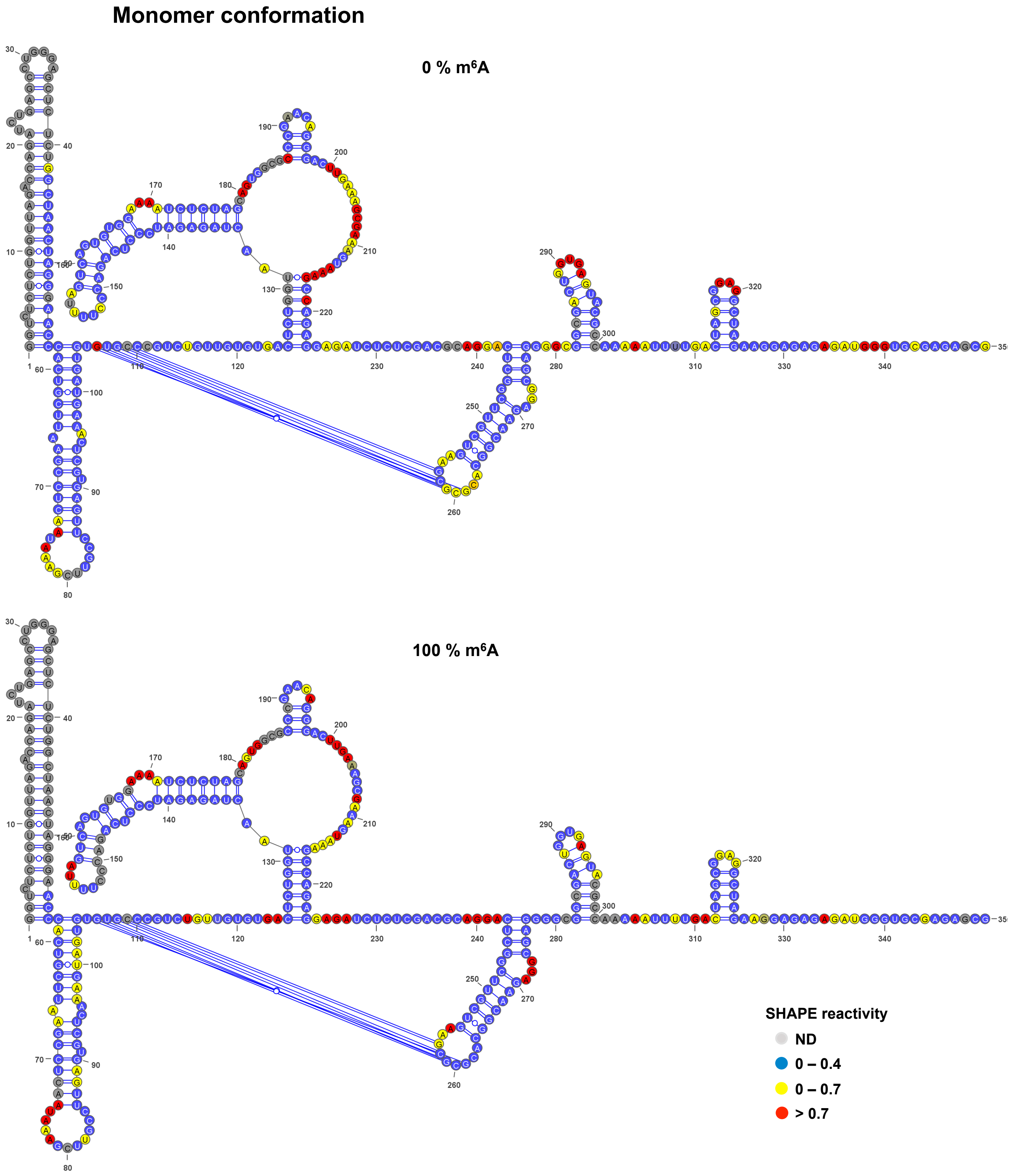


**Supplementary Figure 1. The presence of m$^6$A alters the folding and dimerization of the 5´-UTR of the full-length RNA *in vitro.*** Normalized SHAPE reactivities obtained for the 0% m$^6$A 5'-UTR (upper) and the 100% m$^6$A 5'-UTR (lower).

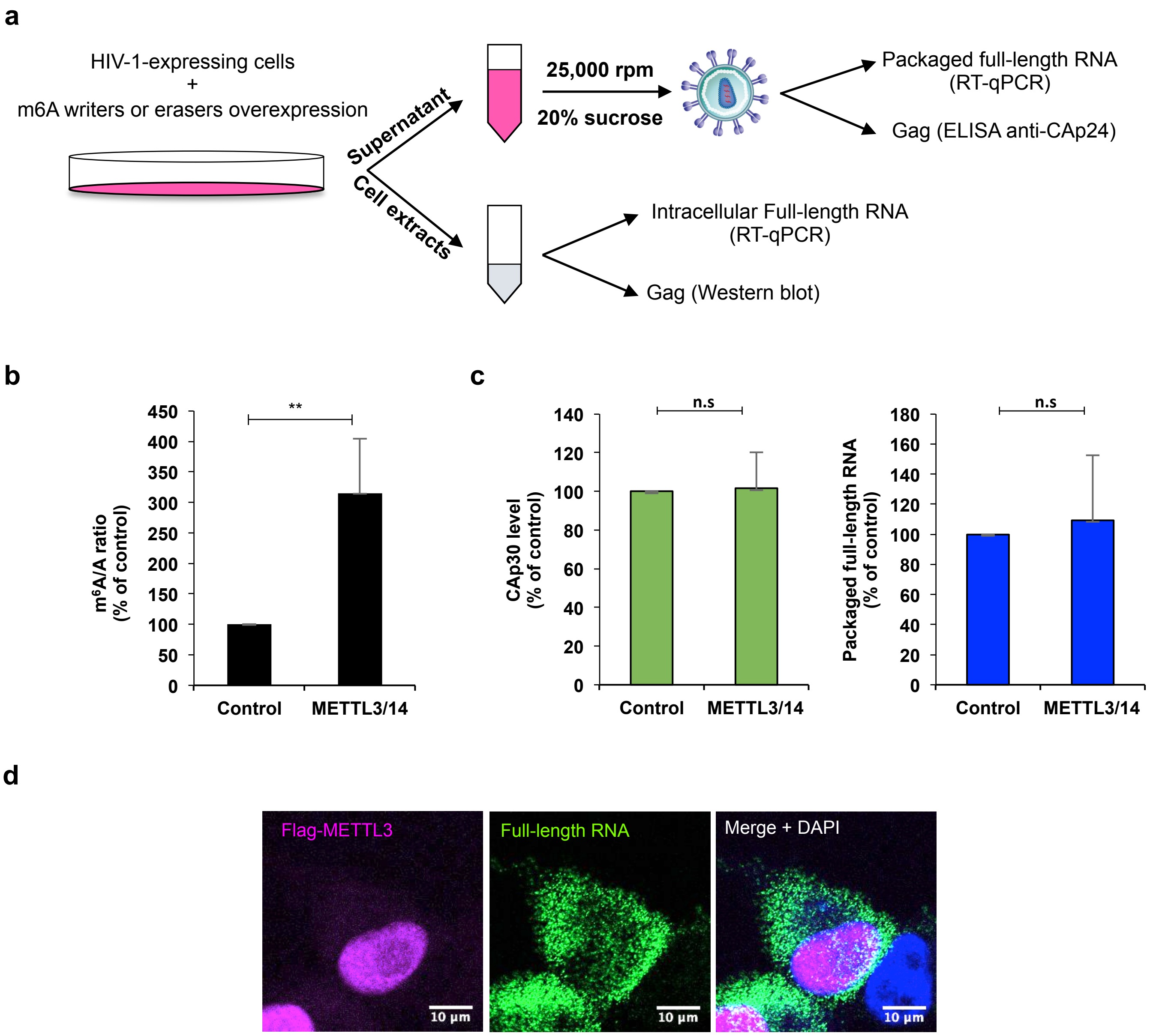


**Supplementary Figure 2. The presence of m$^6$A within the full-length RNA favors Gag synthesis but interferes with packaging.** a, Scheme of the experimental model. Briefly, HEK293T cells were transfected with proviral vector, pCMV-VSVg and the m$^6$A writers or eraser expressing vectors. Supernatant and cell extracts are recovered at 24 hpt. The supernatants are filtered by passing through a 0.22 μm filter and the viral particles are purified by ultracentrifugation at 25.000 rpm for 2 hours at 4ºC in a 20% sucrose cushion. The purified viral particles are used to perform an anti-CAp24 ELISA and RT-qPCR to determinate the level of full-length RNA from CAp24 equivalents. In parallel, the cell extracts from producer cells are used to detect Gag and the overexpressed proteins by Western blot and for RNA extraction to determinate the level of the full-length RNA by RT-qPCR. b, HEK293T cells were transfected with pNL4.3 and pCMV-VSVg together with pCDNA-Flag-METTL3 and pCDNA-Flag-METTL14 or pCDNA-d2EGFP as a control. At 24 hpt cells extracts were used to perform RNA extraction followed by an immunoprecipitation using an anti-m$^6$A antibody (m$^6$A-RIP as described in Methods). The full-length RNA from the input ("A" fraction) and from the immunoprecipitated material ("m$^6$A" fraction) was quantified by RT-qPCR. The m$^6$A/A ratio was normalized to the control (arbitrary set to 100%) and presented as the mean +/-SD of three independent experiments (***$P<0.01$*, *t*-test). c, HEK293T cells were transfected with the MLV proviral DNA pNCAC together with pCDNA-Flag-muMETTL3 and pCDNA-Flag-muMETTL14 or pCDNA-d2EGFP as a control. At 24 hpt the supernatant was filtered and viral particles were purified by ultracentrifugation. Purified viral particles were analyzed by Western blot using and anti-CAp30 antibody and used to perform a densitometric analysis using ImageJ. Additionally, the viral particles were used for RNA extraction and the full-length RNA from equivalents of CAp30 was quantified by RT-qPCR. The levels of CAp30 and the packaged MLV full-length RNA were normalized to the control (arbitrary set to 100%) and presented as the mean +/- SD of three independent experiments (n.s; non-significant, *t*-test). d, HeLa cells were co-transfected with pNL4.3, pCMV-VSVg and pCDNA-Flag-METTL3. At 24 hpt, the expression of Flag-METTL3 and the full-length RNA was determined by FISH and immunofluorescence analyses performed in parallel to the ISH-PLA showed in Fig. 2d. Scale bar 10 μm.

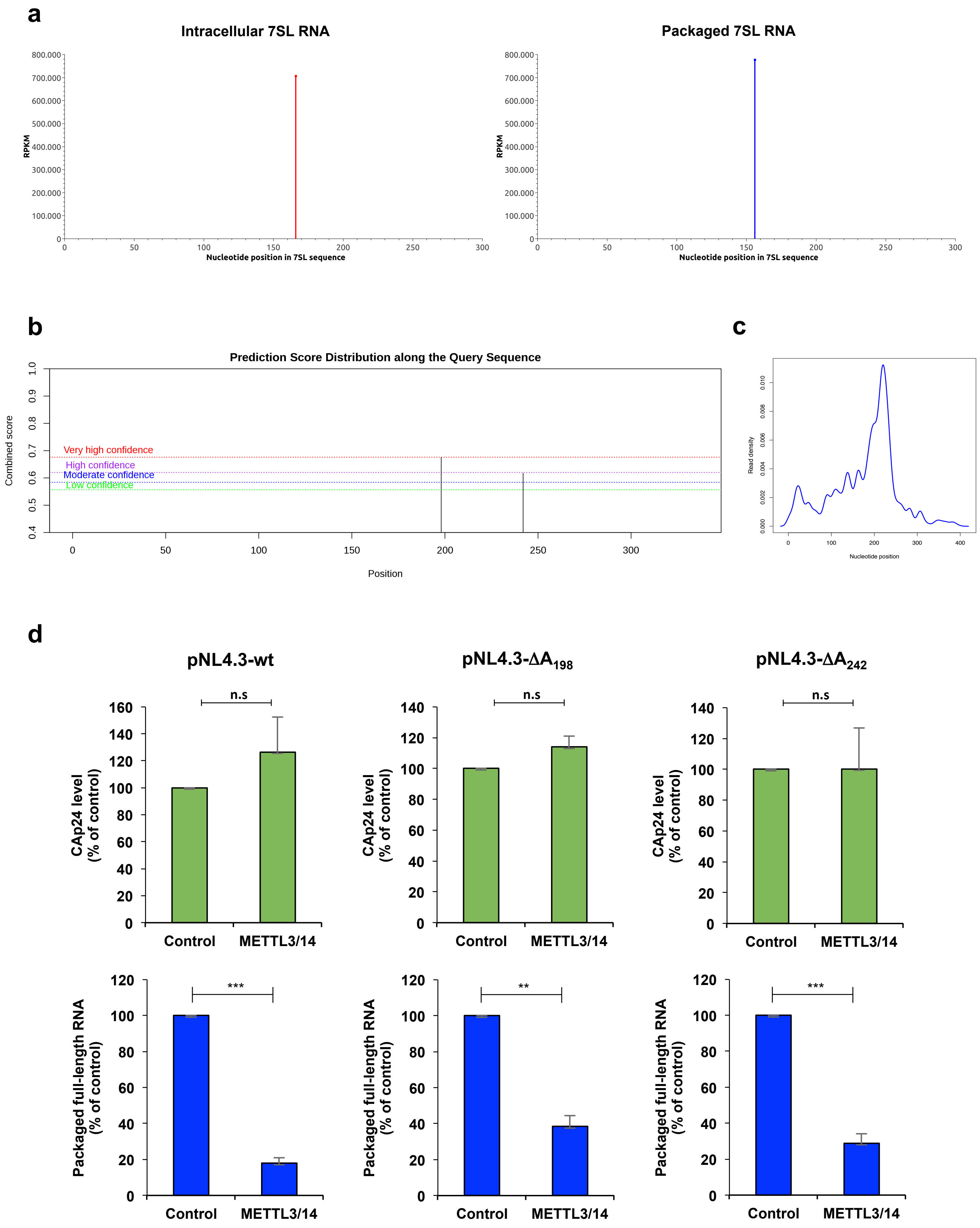

a
Intracellular 7SL RNA
Packaged 7SL RNA
RPKM
Nucleotide position in 7SL sequence
b
Prediction Score Distribution along the Query Sequence
Very high confidence
High confidence
Moderate confidence
Low confidence
Combined score
Position
c
Read density
Nucleotide position
d
pNL4.3-wt
pNL4.3-ΔA198
pNL4.3-ΔA242
n.s
CAp24 level (% of control)
Packaged full-length RNA (% of control)
***
**
Control
METTL3/14

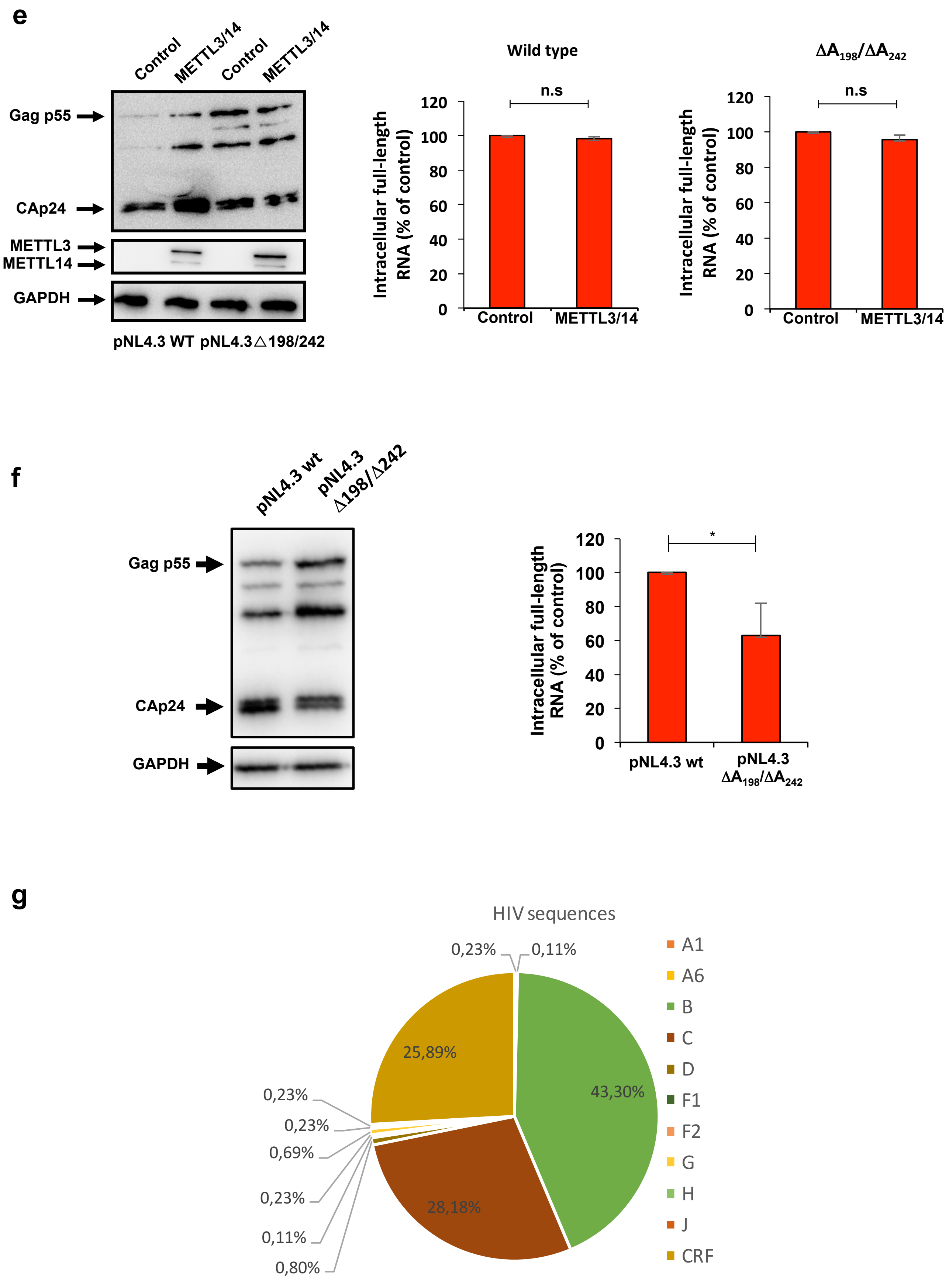
e
Control
METTL3/14
Control
METTL3/14
Gag p55
CAp24
METTL3
METTL14
GAPDH
pNL4.3 WT pNL4.3△198/242
Wild type
n.s
Intracellular full-length RNA (% of control)
120
100
80
60
40
20
0
Control
METTL3/14
ΔA198/ΔA242
n.s
Intracellular full-length RNA (% of control)
Control
METTL3/14
f
pNL4.3 wt
pNL4.3 Δ198/Δ242
Gag p55
CAp24
GAPDH
*
Intracellular full-length RNA (% of control)
120
100
80
60
40
20
0
pNL4.3 wt
pNL4.3 ΔA198/ΔA242
g
HIV sequences
0,23%
0,11%
25,89%
43,30%
28,18%
0,23%
0,23%
0,69%
0,23%
0,11%
0,80%
A1
A6
B
C
D
F1
F2
G
H
J
CRF

**Supplementary Figure 3. Methylation of $A_{198}$ and $A_{242}$ within the 5´-UTR interferes with HIV-1 full-length RNA packaging.** a, Methylation patterns of the host 7SL RNA present within the cells (left) and within HIV-1 particles (right). b, Prediction of possible methylated sites in the 5´-UTR of full length RNA of HIV-1 using the SRAMP software (http://www.cuilab.cn/sramp). This predictor show that $A_{198}$ and $A_{242}$ are two possible methylated residues with very high and high confidence, respectively. c, Zoom of the 5´-UTR methylation peak in intracellular full-length RNA showed in the $m^6A$-seq data of Fig. 3a. d, HEK293T cells were transfected with pNL4.3 wild type (left panel) or pNL4.3-$\Delta A_{198}$ (middle panel) or pNL4.3-$\Delta A_{242}$ (right panel) together with pCMV-VSVg, pCDNA-Flag-METTL3 and pCDNA-Flag-METTL14 or pCDNA-d2EGFP as a control. At 24 hpt, supernatants were filtered and viral particles were purified by ultracentrifugation. Purified viral particles were used to perform an anti-CAp24 ELISA (upper panels) and RNA extraction and RT-qPCR analysis (lower panel). The levels of CAp24 and the packaged full-length RNA (per CAp24 equivalents) were normalized to the control (arbitrary set to 100%) and presented as the mean +/- SD of three independent experiments (**$P<0.01$; ***$P<0,001$, $t$-test). e, HEK293T cells were transfected with pNL4.3 wild type or pNL4.3-$\Delta A_{198}/\Delta A_{242}$ together with pCMV-VSVg, pCDNA-Flag-METTL3 and pCDNA-Flag-METTL14 or pCDNA-d2EGFP as a control. At 24 hpt cells extracts were used to detect Gag, Flag-METTL3 and Flag-METTL14 by Western blot. GAPDH was used as a loading control (left panel). In parallel, cells extracts were used to perform RNA extraction and the full-length RNA was quantified by RT-qPCR (right panel). Intracellular full-length RNA was normalized to the control (arbitrary set to 100%) and presented as the mean +/- SD of three independent experiments (n.s; non-significant, $t$-test). f, HEK293T cells were transfected with pNL4.3 wild type or pNL4.3 $\Delta A_{198}/\Delta A_{242}$ together with pCMV-VSVg. At 24 hpt cells extracts were used to detect Gag, Flag-METTL3 and Flag-METTL14 by Western blot. GAPDH was used as a loading control (left panel). In parallel, cells extracts were used to perform RNA extraction and the full-length RNA was quantified by RT-qPCR (right panel). Intracellular full-length RNA was normalized to the control (arbitrary set to 100%) and presented as the mean +/- SD of three independent experiments (*$P<0.05$, $t$-test). g, Pie chart showing the coverage HIV-1 subtypes sequences used in Fig. 3d.

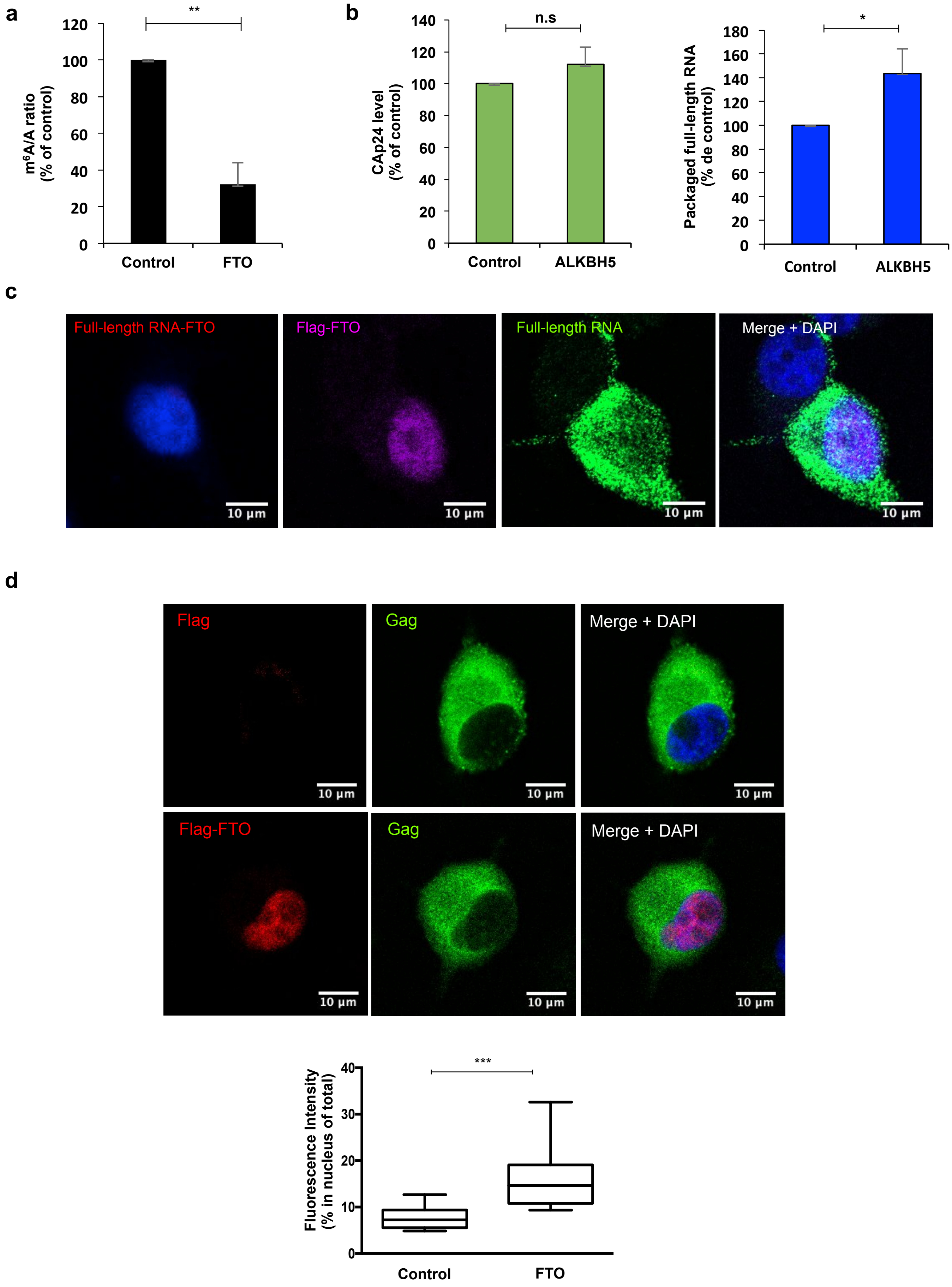
a
**
m6A/A ratio (% of control)
0
20
40
60
80
100
120
Control
FTO
b
n.s
CAp24 level (% of control)
0
20
40
60
80
100
120
140
Control
ALKBH5
*
Packaged full-length RNA (% de control)
0
20
40
60
80
100
120
140
160
180
Control
ALKBH5
c
Full-length RNA-FTO
Flag-FTO
Full-length RNA
Merge + DAPI
10 µm
d
Flag
Gag
Merge + DAPI
Flag-FTO
Gag
Merge + DAPI
10 µm
***
Fluorescence Intensity (% in nucleus of total)
0
10
20
30
40
Control
FTO

**Supplementary Figure 4. Demethylation by a Gag-FTO complex favors HIV-1 full-length RNA packaging.** a, HEK293T cells were transfected with pNL4.3 and pCMV-VSVg together with pCDNA-3XFlag-FTO or pCDNA-d2EGFP as a control. At 24 hpt cells extracts were to perform RNA extraction followed by an immunoprecipitation using an anti-$m^6A$ antibody ($m^6A$-RIP as described in Methods). The full-length RNA from the input ("A" fraction) and from the immunoprecipitated material ("$m^6A$" fraction) was quantified by RT-qPCR. The $m^6A/A$ ratio was normalized to the control (arbitrary set to 100%) and presented as the mean +/-SD of three independent experiments (***$P<0.01$, t*-test). b, HEK293T cells were transfected with pNL4.3 and pCMV-VSVg together with pEGFP-ALKBH5 or pEGFP as a control. At 24 hpt, the supernatant was filtered and ultracentrifuged. Purified viral particles were used to perform an anti-CAp24 ELISA and for RNA extraction and RT-qPCR analysis. The levels of CAp24 and the packaged full-length RNA (per CAp24 equivalents) were normalized to the control (arbitrary set to 100%) and presented as the mean +/- SD of three independent experiments (*$P<0.05$, *t*-test). c, HeLa cells were co-transfected with pNL4.3, pCMV-VSVg and pCDNA-3XFlag-FTO. At 24 hpt, the interaction between the full-length RNA and the Flag-tagged FTO was analyzed by the ISH-PLA protocol described in Methods. The expression of 3XFlag-FTO and the full-length RNA was determined by FISH and immunofluorescence analyses performed in parallel. Scale bar 10 μm. d, HeLa cells were transfected with pNL4.3, pCMV-VSVg and pCDNA-3XFlag-FTO (or pCDNA-Renilla as a control). At 24 hpt, the expression of Flag-FTO and Gag was determined by immunofluorescence analyses. Scale bar 10 μm. The level of Gag in the nucleus (co-localizing with DAPI staining) under control (12 cells) or FTO overexpression (14 cells) was quantified using FIJI/ImageJ (***$P<0,001$*, Mann-Whitney test).
