## Supplementary Methods for "An epitranscriptomic switch at the 5′-UTR controls genome selection during HIV-1 genomic RNA packaging"

**DNA constructs:** The pNL4.3ΔEnv provirus (from here named as pNL4.3) was previously described [1]. The pNL4.3-Δ198, pNL4.3-Δ242 and pNL4.3-Δ198/Δ242 mutant proviruses were obtained by site-directed mutagenesis using pNL4.3 as a template and the primers showed in table 1. pCMV-VSVg was previously described [2]. The pCDNA-Flag-METTL3 and pCDNA-Flag-METTL14 were a gift from Dr. Chuan He (Addgene plasmid #53739) and were previously described [3]. pCDNA-d2EGFP was previously described [1]. The pCDNA-3XFlag-FTO and pEGFP-ALKBH5 were a gift from Dr. Yun-Gui Yang (Beijing Institute of Genomics, Chinese Academy of Sciences) and were previously described [4, 5]. pCDNA-3XFlag-GFP was generated by replacing the FTO cDNA from pCDNA-3XFlag-FTO by the d2EGFP ORF. The pCDNA-Flag-METTL3 and pCDNA-Flag-METTL14 from mouse were a gift from Dr. Jin Crystal Zhao (Sanford Burnham Prebys) and were previously described [6]. The MLV proviral DNA pNCAC was a gift from Dr. Mónica Roth (University of Wisconsin) and was previously described [7]. The pNL4.3-GagStop was previously described [8]. The pCDNA-Gag was previously described [9]. The pBSK-Gag/Pol used for the generation of biotinylated probes was previously described [8].

Table 1: Primers used in mutagenesis reactions

| Primer | Sequence |
|---|---|
| HIV-1Δ198 | Fwd: 5`-TGGCGCCCGAACAGGGCTTGAAAGCGAAAGT-3` |
| | Rev: 5`-ACTTTCGCTTTCAAGCCCTGTTCGGGCGCCA-3` |
| HIV-1Δ242 | Fwd: 5`-ATCTCTCGACGCAGGCTCGGCTTGCTGAAG-3` |
| | Rev: 5`-CTTCAGCAAGCCGAGCCTGCGTCGAGAGAT-3′ |

**Western blot:** Extracts from transfected HEK293T cells were prepared by resuspending cell pellets in lysis buffer (150mM NaCl, 1% NP-40 and 50mM TrisHCl pH 8.0). 30 μg of total protein from cell lysates or MLV viral particles were subjected to 10% SDS-PAGE and transferred to an Amersham Hybond-P 0.45 PVDF membrane (GE Healthcare) for 2 hours, 0.4 A at 4ºC. Membranes were blocked with Blotting-Grade Blocker (BioRad) for 1 hour at room temperature and were incubated with the corresponding primary antibody; a mouse HIV-1 p24 monoclonal antibody diluted to 1/1000 (NIH AIDS Reagents Program, Catalog number 3537), rabbit MLV CAp antibody diluted 1/5000 (kindly provided by Dr. Gloria

Arriagada, UNAB, Chile), a rabbit anti-Flag antibody (Sigma-Aldrich Catalog number 637303) diluted to 1/1000 or a mouse anti-GAPDH diluted 1/5000 (Santa Cruz Biotechnologies Catalog number sc-51905). Upon incubation with the corresponding HRP-conjugated secondary antibody (Jackson ImmunoResearch) diluted to 1/5000, membranes were analyzed with the Pierce® ECL Substrate or the SuperSignal™ West Femto (Thermo Scientific) using a MiniHD9 Western Blot Imaging System scanner (Uvitec Cambridge).

**ELISA:** The anti-CAp24 ELISA (R&D SYSTEMS, catalog number DHP240B) was performed as indicated by the manufacturer. Briefly, viral particles in PBS were mixed with the Calibrator Diluent RD5-26 (diluted 1/4). Then, 100 μl of the Assay Diluent RD1-124 were added in the well coated with a monoclonal antibody specific for HIV-1 Gag p24 and 100 μl of PBS (control) or the sample prepared previously. The wells were incubated for 2 hours at room temperature on a horizontal orbital microplate shaker at 500 rpm, washed three times with Wash Buffer and incubated with 200 μl of HIV-1 Gag p24 Conjugate for 2 hours at room temperature on the shaker. After the incubation, the wells were washed again and incubated with 200 μl of Substrate Solution for 30 minutes at room temperature protected from the light. The reaction was stopped with 50 μl of Stop Solution and the optical density was measured at 450 nm and corrected at 540 nm in a Synergy HTX multi-mode reader (BioTek).

**RNA extraction and RT-qPCR:** RNA extraction and RT-qPCR was performed essentially as recently described [8]. Briefly, cells were washed and recovered with PBS-EDTA 10 mM. Cells in PBS were centrifuged at 5000 rpm for 5 min at 4 ºC cell pellets were resuspended in 200 μl of PBS. Cells or purified viral particles were used to perform RNA extraction by adding 1 ml of TRIzol™ (Thermo Fisher) and 200 μl of chloroform (Merck). The mix was vortexed vigorously and centrifuged at 10.000 rpm for 5 minutes at 4 ºC. The aqueous phase was recovered, incubated for 5 minutes with an equal volume of 2-propanol (Merck) and centrifuged at 12.000 rpm for 20 minutes at 4 ºC. The RNA pellet was washed with ETOH 70% (Merck) and resuspended in ultrapure water. RNAs obtained were treated with TURBO DNA-free kit (Thermo Fisher) for 30 minutes at 37 ºC as indicated by the manufacturer. RNAs were reverse-transcribed using the High Capacity RNA-to-cDNA Master Mix (Thermo Fisher) as indicated by the manufacturer. For intracellular RNAs, 300 ng were used to perform the reverse transcription. For RNA obtained from viral particles RNA, the amount of

RNA used for the reverse transcription reaction was normalized to the CAp24 levels obtained by ELISA using 300 ng of RNA from the control condition. For the qPCR, a 25 µl reaction mix was prepared with 5 µl of template cDNAs (previously diluted to 1/10), 12.5 µl of Brilliant II SYBR® Green QPCR Master Mix (Agilent Technologies), 0.4 µl of sense and antisense primers (stored in a mix a 10 mM) and 7.1 µl of water. The reaction mix was subjected to amplification using the AriaMx Real-Time PCR System (Agilent Technologies). The GAPDH housekeeping gene was amplified in parallel to serve as a control reference. Relative copy numbers of HIV-1 and MLV full-length RNA were compared to GAPDH using $x^{-\Delta Ct}$ (where *x* correspond to the experimentally calculated amplification efficiency of each primer couple).

Table 2: Primers used in qPCR reactions

| Primer | Sequence |
|---|---|
| HIV-1 FL RNA | Fwd: 5`- GCAGTGGCGCCCGAACAGG -3` |
| | Rev: 5`- TTTTTGGCGTACTCACCAGTC -3` |
| MLV FL RNA | Fwd: 5`- TGATCTTAACCTGGGTGATGAG-3` |
| | Rev: 5`- ATCGCTCACAACCAGTCGG-3′ |
| GAPDH | Fwd: 5`- AGCCACATCGCTCAGACAC-3` |
| | Rev: 5`- GCCCAATACGACCAAATCC-3` |

**Methylated RNA immunoprecipitation (MeRIP):** RNA extraction from overexpressing cells (METTL3/14 or FTO) or from cells expressing wild type or GagStop proviral DNA was performed from cells extracts as described above. For MeRIP, 30 µg of total RNA diluted in 380 µl of water was denatured at 70 ºC for 5 min and placed on ice for 3 min. The denatured RNA was mixed with 1 µl of rRNasin® (Promega), 5 µl VRC, 100 µl of IP Buffer 5X (50 mM Tris-Hcl pH7.4, 750 mM NaCl and 0.5% NP-40) and 2,5 µl of rabbit polyclonal anti-$m^6A$ (1 mg/mL; Abcam #ab151230) and incubated for 2 hours at 4 ºC with head-over-tail rotation. At the same time, 600 µg of Dynabeads™ Protein A magnetic beads (20 µl; Thermo Fisher) were washed in 1 ml of IP Buffer 1X with 1 µl of VRC and were incubated with 500 µl of Buffer IP 1X with 0.5 mg/ml of BSA for 2 hours at 4 ºC with head-over-tail rotation. Then, beads were washed with 500 µl of IP Buffer 1X and added to the RNA/anti-$m^6A$ antibody mix. The RNA-beads mix was incubated for 2 hours at 4ºC in head-over-tail rotation. After incubation, the RNA-beads mix was washed twice with 500 µl IP Buffer 1X.

Bound RNA was eluted with 100 µl of Elution Buffer (5mM Tris-HCl, 1mM EDTA and 0.05% SDS) and 1 µl of Proteinase K (New England BioLabs) and incubated for 1.5 hour at 50ºC. RNA was extracted from supernatant using TRIzol™ (Thermo Fisher). The RNA recovered was precipitated with 10 mM $MgCl_2$, 20 µg glycogen (Thermo Fisher) and 2.5 volumes of ETOH 100% overnight at -20 ºC and then washed with ETOH 70%. Equal amounts of RNA from input and immunoprecipitation were used for RT-qPCR. The full-length RNA from the immunoprecipitated material ("$m^6A$" fraction) was normalized to full-length RNA from the input ("A" fraction).

**Prediction of methylated residues:** Prediction of methylated adenosines within the 5′-UTR of the NL4.3 strain was perfomed with the SRAMP software available at www.cuilab.cn/sramp considering secondary RNA structure [10].

**Fluorescent in situ hybridization, immunofluorescence and confocal microscopy:** Specific probes against the HIV-1 full-length RNA were generated by *in vitro* transcription using the pBSK-Gag/Pol vector and digoxin-11-UTP (Roche) as we previously described [1]. The generated 5-kb transcript complementary to the Gag/Pol region of the full-length RNA was fragmented using the RNA fragmentation buffer (Thermo Scientific) in order to obtain probes of 100–200 nt in length following the supplier′s instructions. Fragmented probes were purified using the Agencourt AMPure XP magnetic beads (Beckman Coulter). RNA FISH was carried out as we recently described [1]. Briefy, HeLa cells were cultured in a 12-well plate with coverslips and maintained as described in Methods. Cells were transfected with 0.3 µg of pNL4.3, 0.3 µg of pCMV-VSVg and 1 µg of the corresponding expression vectors as indicated in Methods. At 24 hpt, cells were washed twice with 1× PBS and fixed for 10 min at room temperature with 4% paraformaldehyde. Cells were subsequently permeabilized for 10 min at room temperature with 0.2% Triton X-100 and hybridized overnight at 37 °C in 200 µl of hybridization mix (10% dextran sulfate, 2 mM VRC, 0.02% RNase-free BSA, 50% formamide, 300 µg tRNA and 120 ng of 11-digoxigenin-UTP probes) in a humid chamber. Cells were washed with 0.2X SSC/50% formamide during 30 min at 50ºC and then incubated three times with antibody dilution buffer (2X SSC, 8% formamide, 2 mM vanadyl-ribonucleoside complex, 0.02% RNase-free BSA). Sheep anti-digoxin (Roche), mouse HIV-1 anti-p24 (NIH AIDS Reagents Program, Catalog number 3537) and Rabbit anti-Flag (Sigma) primary antibodies diluted to 1/100 in antibody dilution buffer were added for 2 h at room temperature. After three washes with antibody dilution buffer, cells were incubated for 90 min at room temperature with anti-Sheep Alexa 488, anti-mouse Alexa 594 and anti-rabbit Alexa 647 antibodies (Molecular Probes) diluted at 1/500. Cells were washed three times in wash buffer (2X SSC, 8% formamide, 2 mM vanadyl-ribonucleoside complex), twice with 1X PBS, incubated with DAPI (0.3 μg/ml in PBS) (Life Technologies) for 1 min at room temperature, washed three times with 1X PBS, three times with water and mounted with Fluoromount™ (Life Technologies). Images were obtained with a Zeiss LSM 800 Confocal Microscope (Zeiss) and processed using FIJI/ImageJ (NIH).

**Proximity ligation (PLA):** PLA was carried out using the DUOLINK II In Situ kit (Sigma-Aldrich) and PLA probe anti-mouse minus and PLA probe anti-rabbit plus (Sigma-Aldrich) as we have previously described [1]. Briefly, HeLa cells transfected with pNL4.3 and pCDNA-3XFlag-FTO for 24 hours were fixed and pre-incubated with blocking agent for 30 min at room temperature. Primary antibodies were added at a dilution of 1/100 (mouse anti-HIV-1 p24 monoclonal antibody) and 1/200 (rabbit anti-Flag, Sigma-Aldrich) in 40 μl DUOLINK antibody diluent and incubated at 37 ºC for 1 h. Samples were washed three times with PBS for 5 min each and secondary antibodies (DUOLINK anti-rabbit PLA-plus probe and DUOLINK anti-mouse PLA-minus probes) were added and incubated at 37 ºC for 1 h. Ligation and amplification reactions were performed following the same protocol described in [1]. Samples were incubated with DAPI (0.3 μg/ml in PBS) (Life Technologies) for 1 min at room temperature and coverslips were washed three times with PBS, three times with water and mounted with Fluoromount™ (Sigma Aldrich). Images were obtained with a Zeiss LSM 800 Confocal Microscope (Zeiss) and processed using FIJI/ImageJ (NIH).

**In situ hybridization coupled to PLA (ISH-PLA):** The ISH-PLA protocol was developed by mixing the RNA-FISH and PLA protocols described above. Briefly, HeLa cells transfected with pNL4.3 and pCDNA-Flag-METTL3 or pCDNA-3XFlag-FTO growing on coverslips were fixed, permeabilized for 10 min at RT with 0.2% Triton X-100 and hybridized overnight at 37 ºC in 200 μl of hybridization mix (10% dextran sulphate, 2 mM vanadyl–ribonucleoside complex, 0.02% RNase-free bovine serum albumin, 50% formamide, 300 μg of tRNA and 120 ng of 11-digoxigenin-UTP probes) in a humid chamber. Cells were washed with 0.2X SSC/50% formamide during 30 min at 50°C and incubated with blocking agent for 30 min to room temperature. Cells were then incubated three times with antibody dilution buffer (2X

SSC, 8% formamide, 2 mM vanadyl–ribonucleoside complex and 0.02% RNase-free bovine serum albumin). Mouse anti-digoxin and rabbit anti-flag primary antibodies diluted to 1/100 in antibody dilution buffer were added for 2 h at room temperature. After three washes with antibody dilution buffer and two washes with PBS for 5 min each, the secondary antibodies (DUOLINK anti-rabbit PLA-plus probe, DUOLINK anti-mouse PLA minus probe) were added and incubated at 37 ºC for 1 h. The ligation and amplification reactions were performed following the same protocol described above. Thereafter, coverslips were incubated with a solution of DAPI (0.3 μg/ml in PBS) (Life Technologies) for 1 min at room temperature, washed three times with PBS, three times with water and mounted with Fluoromount™ (Sigma Aldrich). Images were obtained with a Zeiss LSM 800 Confocal Microscope (Zeiss) and processed using FIJI/ImageJ (NIH).

**Adenosine conservation analysis:** For the adenosine conservation analysis, all the HIV-1 sequences present in HIV database were aligned with *HIV Premade Aligments*, ALL GENOME options, processed and classified with an *in house* script. Motifs surrounding $A_{198}$ and $A_{242}$ were analyzed by *WebLogo* v2.8.2 (https://weblogo.berkeley.edu/logo.cgi).

### Monomer conformation

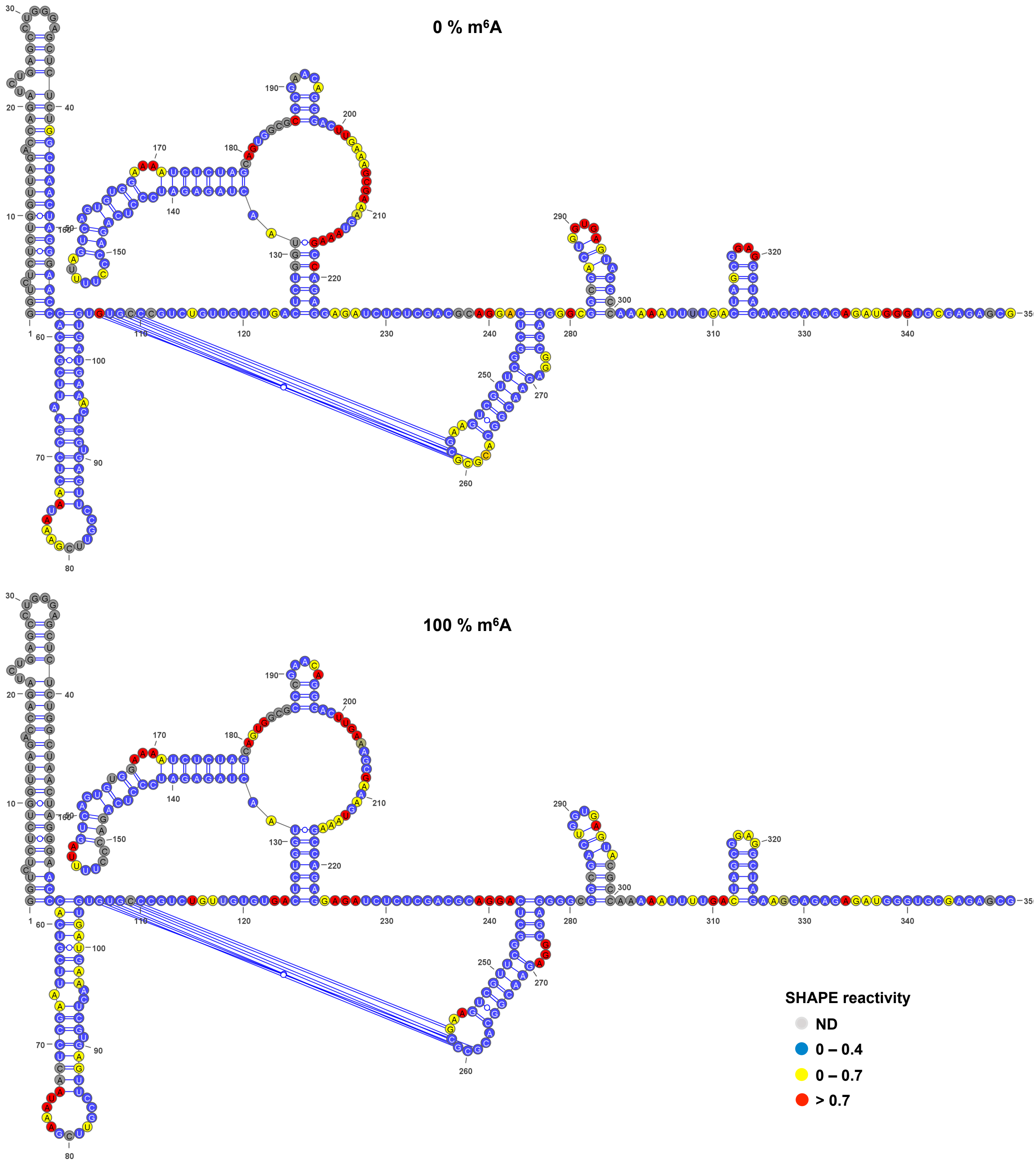


**Supplementary Figure 1. The presence of m$^6$A alters the folding and dimerization of the 5´-UTR of the full-length RNA *in vitro.*** Normalized SHAPE reactivities obtained for the 0% m$^6$A 5'-UTR (upper) and the 100% m$^6$A 5'-UTR (lower).

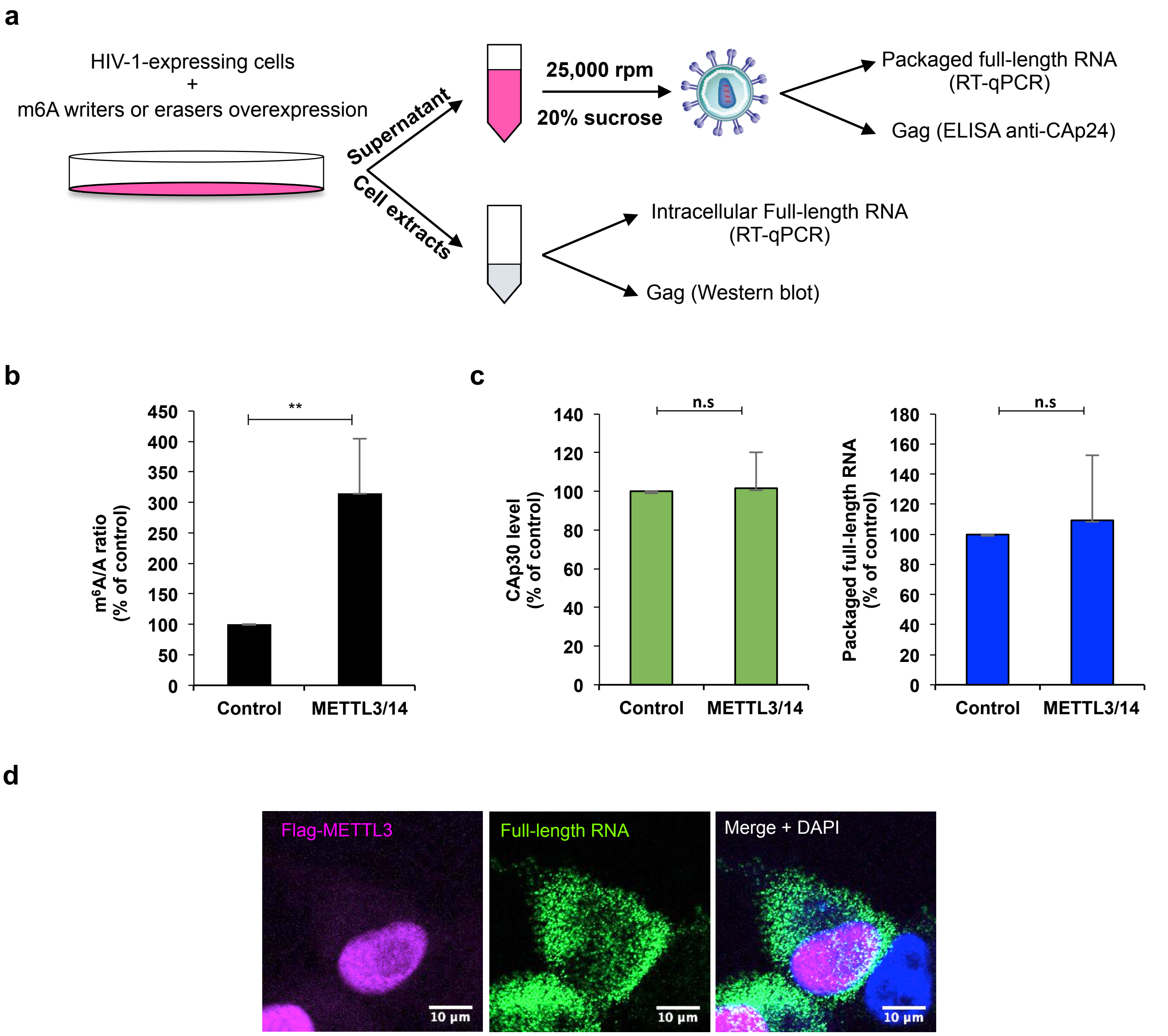


**Supplementary Figure 2. The presence of m⁶A within the full-length RNA favors Gag synthesis but interferes with packaging.** a, Scheme of the experimental model. Briefly, HEK293T cells were transfected with proviral vector, pCMV-VSVg and the m⁶A writers or eraser expressing vectors. Supernatant and cell extracts are recovered at 24 hpt. The supernatants are filtered by passing through a 0.22 µm filter and the viral particles are purified by ultracentrifugation at 25.000 rpm for 2 hours at 4ºC in a 20% sucrose cushion. The purified viral particles are used to perform an anti-CAp24 ELISA and RT-qPCR to determinate the level of full-length RNA from CAp24 equivalents. In parallel, the cell extracts from producer cells are used to detect Gag and the overexpressed proteins by Western blot and for RNA extraction to determinate the level of the full-length RNA by RT-qPCR. b, HEK293T cells were transfected with pNL4.3 and pCMV-VSVg together with pCDNA-Flag-METTL3 and pCDNA-Flag-METTL14 or pCDNA-d2EGFP as a control. At 24 hpt cells extracts were used to perform RNA extraction followed by an immunoprecipitation using an anti-m⁶A antibody (m⁶A-RIP as described in Methods). The full-length RNA from the input ("A" fraction) and from the immunoprecipitated material ("m⁶A" fraction) was quantified by RT-qPCR. The m⁶A/A ratio was normalized to the control (arbitrary set to 100%) and presented as the mean +/-SD of three independent experiments (***P<0.01,* *t*-test). c, HEK293T cells were transfected with the MLV proviral DNA pNCAC together with pCDNA-Flag-muMETTL3 and pCDNA-Flag-muMETTL14 or pCDNA-d2EGFP as a control. At 24 hpt the supernatant was filtered and viral particles were purified by ultracentrifugation. Purified viral particles were analyzed by Western blot using and anti-CAp30 antibody and used to perform a densitometric analysis using ImageJ. Additionally, the viral particles were used for RNA extraction and the full-length RNA from equivalents of CAp30 was quantified by RT-qPCR. The levels of CAp30 and the packaged MLV full-length RNA were normalized to the control (arbitrary set to 100%) and presented as the mean +/- SD of three independent experiments (n.s; non-significant, *t*-test). d, HeLa cells were co-transfected with pNL4.3, pCMV-VSVg and pCDNA-Flag-METTL3. At 24 hpt, the expression of Flag-METTL3 and the full-length RNA was determined by FISH and immunofluorescence analyses performed in parallel to the ISH-PLA showed in Fig. 2d. Scale bar 10 µm.

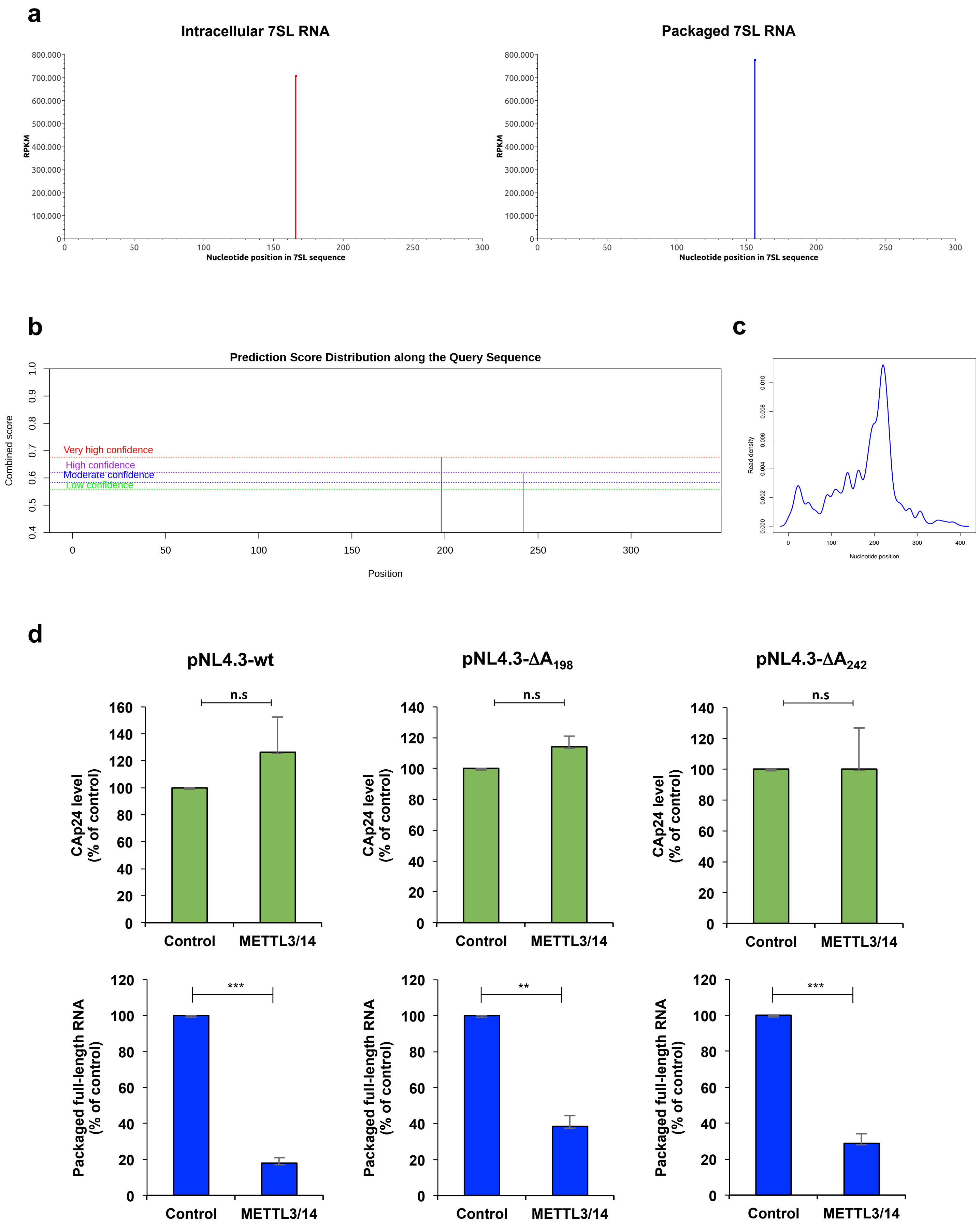

a
Intracellular 7SL RNA
Packaged 7SL RNA
RPKM
Nucleotide position in 7SL sequence
b
Prediction Score Distribution along the Query Sequence
Combined score
Very high confidence
High confidence
Moderate confidence
Low confidence
Position
c
Read density
Nucleotide position
d
pNL4.3-wt
pNL4.3-ΔA198
pNL4.3-ΔA242
n.s
CAp24 level (% of control)
Packaged full-length RNA (% of control)
***
**
Control
METTL3/14

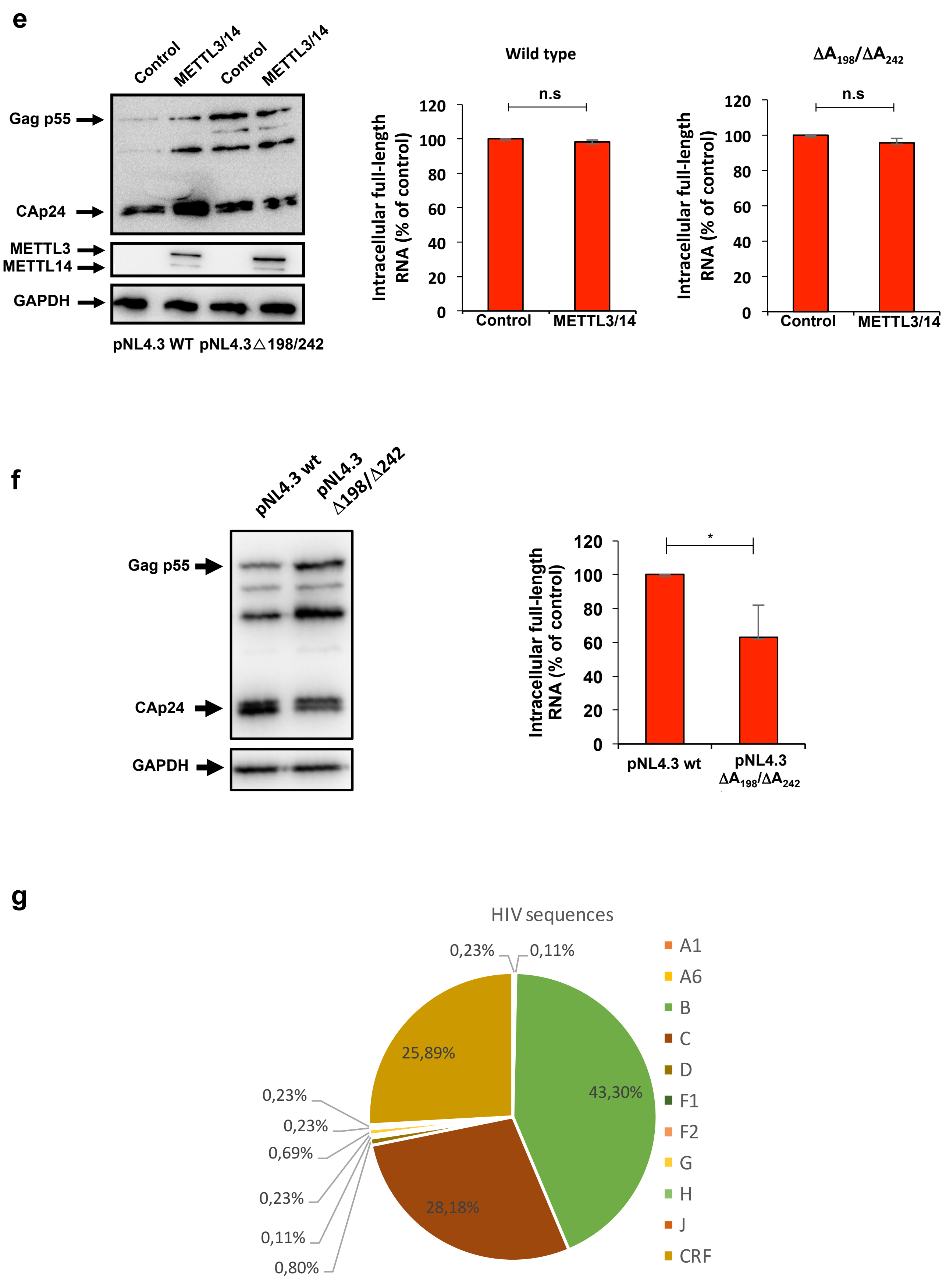
e
Control
METTL3/14
Control
METTL3/14
Gag p55
CAp24
METTL3
METTL14
GAPDH
pNL4.3 WT pNL4.3△198/242
Wild type
$\Delta A_{198}/\Delta A_{242}$
n.s
n.s
Intracellular full-length RNA (% of control)
120
100
80
60
40
20
0
Control
METTL3/14
f
pNL4.3 wt
pNL4.3 Δ198/Δ242
Gag p55
CAp24
GAPDH
*
Intracellular full-length RNA (% of control)
pNL4.3 wt
pNL4.3 $\Delta A_{198}/\Delta A_{242}$
g
HIV sequences
0,23%
0,11%
25,89%
43,30%
28,18%
0,23%
0,23%
0,69%
0,23%
0,11%
0,80%
A1
A6
B
C
D
F1
F2
G
H
J
CRF

**Supplementary Figure 3. Methylation of $A_{198}$ and $A_{242}$ within the 5´-UTR interferes with HIV-1 full-length RNA packaging.** a, Methylation patterns of the host 7SL RNA present within the cells (left) and within HIV-1 particles (right). b, Prediction of possible methylated sites in the 5´-UTR of full length RNA of HIV-1 using the SRAMP software (http://www.cuilab.cn/sramp). This predictor show that $A_{198}$ and $A_{242}$ are two possible methylated residues with very high and high confidence, respectively. c, Zoom of the 5´-UTR methylation peak in intracellular full-length RNA showed in the $m^6A$-seq data of Fig. 3a. d, HEK293T cells were transfected with pNL4.3 wild type (left panel) or pNL4.3-$\Delta A_{198}$ (middle panel) or pNL4.3-$\Delta A_{242}$ (right panel) together with pCMV-VSVg, pCDNA-Flag-METTL3 and pCDNA-Flag-METTL14 or pCDNA-d2EGFP as a control. At 24 hpt, supernatants were filtered and viral particles were purified by ultracentrifugation. Purified viral particles were used to perform an anti-CAp24 ELISA (upper panels) and RNA extraction and RT-qPCR analysis (lower panel). The levels of CAp24 and the packaged full-length RNA (per CAp24 equivalents) were normalized to the control (arbitrary set to 100%) and presented as the mean +/- SD of three independent experiments (***P<0.01*; ****P<0,001, t*-test). e, HEK293T cells were transfected with pNL4.3 wild type or pNL4.3-$\Delta A_{198}/\Delta A_{242}$ together with pCMV-VSVg, pCDNA-Flag-METTL3 and pCDNA-Flag-METTL14 or pCDNA-d2EGFP as a control. At 24 hpt cells extracts were used to detect Gag, Flag-METTL3 and Flag-METTL14 by Western blot. GAPDH was used as a loading control (left panel). In parallel, cells extracts were used to perform RNA extraction and the full-length RNA was quantified by RT-qPCR (right panel). Intracellular full-length RNA was normalized to the control (arbitrary set to 100%) and presented as the mean +/- SD of three independent experiments (n.s; non-significant, *t*-test). f, HEK293T cells were transfected with pNL4.3 wild type or pNL4.3 $\Delta A_{198}/\Delta A_{242}$ together with pCMV-VSVg. At 24 hpt cells extracts were used to detect Gag, Flag-METTL3 and Flag-METTL14 by Western blot. GAPDH was used as a loading control (left panel). In parallel, cells extracts were used to perform RNA extraction and the full-length RNA was quantified by RT-qPCR (right panel). Intracellular full-length RNA was normalized to the control (arbitrary set to 100%) and presented as the mean +/- SD of three independent experiments (**P<0.05, t*-test). g, Pie chart showing the coverage HIV-1 subtypes sequences used in Fig. 3d.

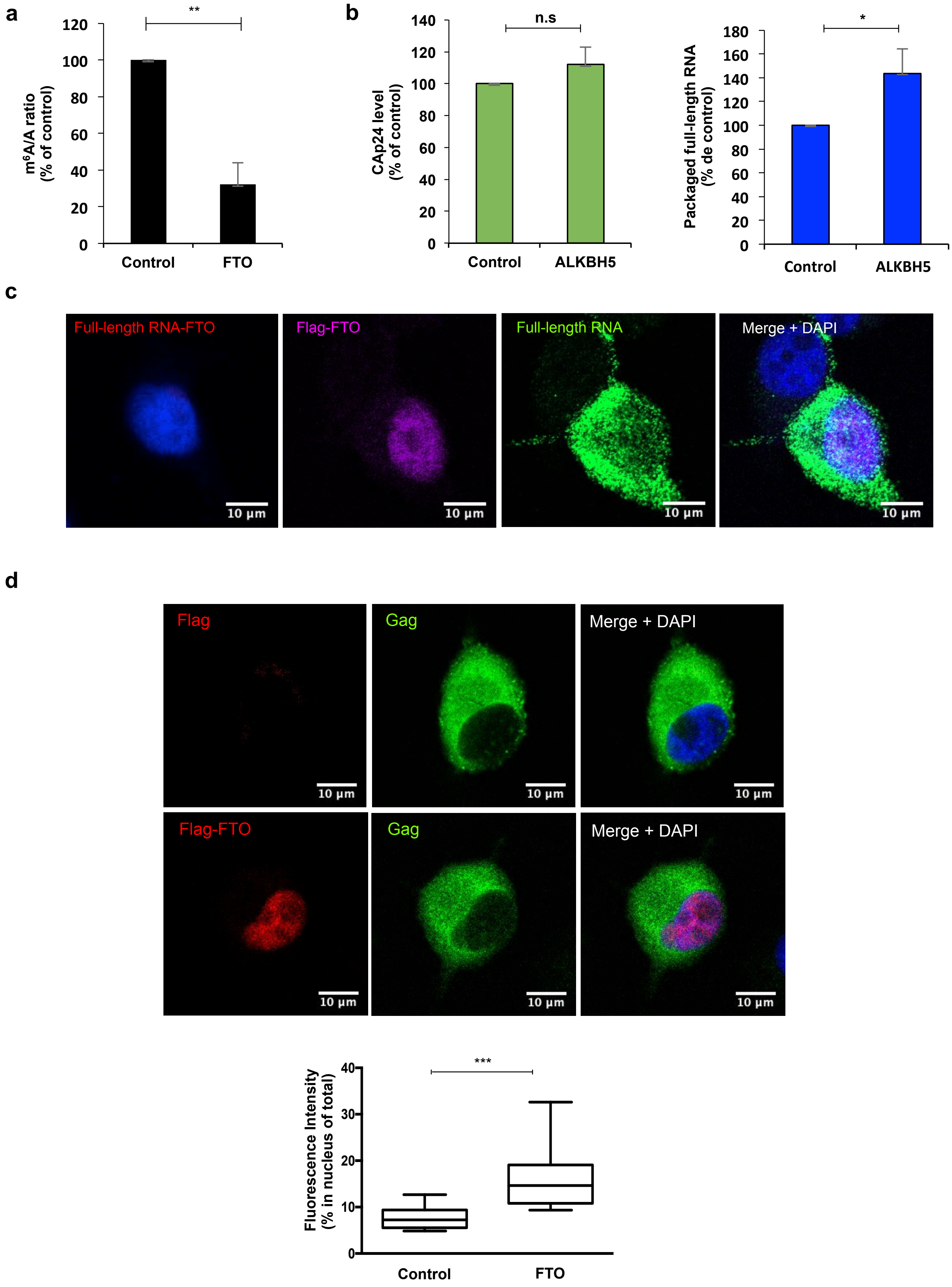
a
**
m6A/A ratio (% of control)
0
20
40
60
80
100
120
Control
FTO
b
n.s
CAp24 level (% of control)
0
20
40
60
80
100
120
140
Control
ALKBH5
*
Packaged full-length RNA (% de control)
0
20
40
60
80
100
120
140
160
180
Control
ALKBH5
c
Full-length RNA-FTO
Flag-FTO
Full-length RNA
Merge + DAPI
10 µm
d
Flag
Gag
Merge + DAPI
Flag-FTO
Gag
Merge + DAPI
10 µm
***
Fluorescence Intensity (% in nucleus of total)
0
10
20
30
40
Control
FTO

**Supplementary Figure 4. Demethylation by a Gag-FTO complex favors HIV-1 full-length RNA packaging.** a, HEK293T cells were transfected with pNL4.3 and pCMV-VSVg together with pCDNA-3XFlag-FTO or pCDNA-d2EGFP as a control. At 24 hpt cells extracts were to perform RNA extraction followed by an immunoprecipitation using an anti-$m^6A$ antibody ($m^6A$-RIP as described in Methods). The full-length RNA from the input ("A" fraction) and from the immunoprecipitated material ("$m^6A$" fraction) was quantified by RT-qPCR. The $m^6A$/A ratio was normalized to the control (arbitrary set to 100%) and presented as the mean +/-SD of three independent experiments (***P<0.01, t*-test). b, HEK293T cells were transfected with pNL4.3 and pCMV-VSVg together with pEGFP-ALKBH5 or pEGFP as a control. At 24 hpt, the supernatant was filtered and ultracentrifuged. Purified viral particles were used to perform an anti-CAp24 ELISA and for RNA extraction and RT-qPCR analysis. The levels of CAp24 and the packaged full-length RNA (per CAp24 equivalents) were normalized to the control (arbitrary set to 100%) and presented as the mean +/- SD of three independent experiments (*$P<0.05$, *t*-test). c, HeLa cells were co-transfected with pNL4.3, pCMV-VSVg and pCDNA-3XFlag-FTO. At 24 hpt, the interaction between the full-length RNA and the Flag-tagged FTO was analyzed by the ISH-PLA protocol described in Methods. The expression of 3XFlag-FTO and the full-length RNA was determined by FISH and immunofluorescence analyses performed in parallel. Scale bar 10 μm. d, HeLa cells were transfected with pNL4.3, pCMV-VSVg and pCDNA-3XFlag-FTO (or pCDNA-Renilla as a control). At 24 hpt, the expression of Flag-FTO and Gag was determined by immunofluorescence analyses. Scale bar 10 μm. The level of Gag in the nucleus (co-localizing with DAPI staining) under control (12 cells) or FTO overexpression (14 cells) was quantified using FIJI/ImageJ (****P<0,001,* Mann-Whitney test).
